# Diet-induced loss of adipose Hexokinase 2 triggers hyperglycemia

**DOI:** 10.1101/2019.12.28.887794

**Authors:** Mitsugu Shimobayashi, Sunil Shetty, Irina C. Frei, Bettina K. Wölnerhanssen, Diana Weissenberger, Nikolaus Dietz, Amandine Thomas, Danilo Ritz, Anne Christin Meyer-Gerspach, Timm Maier, Nissim Hay, Ralph Peterli, Nicolas Rohner, Michael N. Hall

## Abstract

Chronically high blood glucose (hyperglycemia) leads to diabetes, fatty liver disease, and cardiovascular disease. Obesity is a major risk factor for hyperglycemia, but the underlying mechanism is unknown. Here we show that a high fat diet (HFD) in mice causes early loss of expression of the glycolytic enzyme Hexokinase 2 (HK2) specifically in adipose tissue. Adipose-specific knockout of *Hk2* caused enhanced gluconeogenesis and lipogenesis in liver, a condition known as selective insulin resistance, leading to glucose intolerance. Furthermore, we observed reduced hexokinase activity in adipose tissue of obese and diabetic patients, and identified a loss-of-function mutation in the *hk2* gene of naturally hyperglycemic Mexican cavefish. Mechanistically, HFD in mice led to loss of HK2 by inhibiting translation of *Hk2* mRNA. Our findings identify adipose HK2 as a critical mediator of systemic glucose homeostasis, and suggest that obesity-induced loss of adipose HK2 is an evolutionarily conserved, non-cell-autonomous mechanism for the development of hyperglycemia.

**One Sentence Summary:** Loss of the glycolytic enzyme Hexokinase 2 in adipose tissue is a mechanism underlying high blood glucose levels.

## Main text

Vertebrates mediate glucose homeostasis by regulating glucose production and uptake in specific tissues (*1, 2*). High blood glucose stimulates pancreatic beta cells to secrete the hormone insulin which in turn promotes glucose uptake in skeletal muscle and adipose tissue and inhibits glucose production in liver. Although its contribution to glucose clearance is minor (*3, 4*), adipose tissue plays a particularly important role in systemic glucose homeostasis (*5, 6*). Adipose-specific knockout of insulin signaling components, such as the insulin receptor, mTORC2, and AKT, results in local and systemic insulin insensitivity (*7–13*). However, we and others recently reported that diet-induced obesity in mice causes hyperglycemia despite normal insulin signaling in adipose tissue (*14, 15*). These observations raise the question, how does obesity cause insulin insensitivity and hyperglycemia without affecting the insulin signaling pathway? In other words, how does diet induce diabetes?

To determine the molecular basis of diet-induced insulin insensitivity in adipocytes and hyperglycemia in mice, we performed an unbiased proteomic analysis on visceral white adipose tissue (vWAT) isolated from C57BL/6JRj wild-type mice fed an HFD or normal diet (ND) for 4 weeks (**Fig. S1A-C**). We detected and quantified 6294 proteins of which 52 and 67 were up- and down-regulated, respectively, in vWAT of HFD-fed mice (**Table S1**). The glycolytic enzyme Hexokinase 2 (HK2), expressed in adipose tissue and muscle, was among the proteins significantly down-regulated in vWAT (**Fig. 1A**). HK2 phosphorylates glucose to generate glucose-6-phosphate (G6P), the rate-limiting step in glycolysis, in an insulin-stimulated manner. We focused on HK2 due to efflux of non-phosphorylated glucose and thus a potential link of HK2 loss to hyperglycemia. HK2 is the most abundant (~80%) of the three hexokinase isoforms expressed in vWAT but the only one down-regulated upon HFD (**Fig. 1A** and **Fig. S2**). Loss of HK2 was confirmed in vWAT and demonstrated in subcutaneous WAT (sWAT) and brown adipose tissue (BAT) of 4-week HFD mice by immunoblotting (**Fig. 1B**). Consistent with reduced HK2 expression, hexokinase activity was decreased in vWAT and sWAT (**Fig. S3)**. A longitudinal study of HFD-fed mice revealed that HK2 down-regulation in vWAT and sWAT occurred already at one week of HFD and correlated with hyperglycemia (**Fig. S4A-E**). HK2 expression was unchanged in skeletal muscle (**Fig. 1B**), indicating that HFD decreases HK2 expression specifically in adipose tissue. Consistent with a previous study (*14*), expression of the glucose transporter GLUT4 was not affected in vWAT of 4-week HFD mice (**Fig. 1A**). GLUT4 was down-regulated in sWAT after ≥2 weeks of HFD (**Fig. S4E**). Confirming earlier observations (*14, 15*), insulin signaling, as determined by AKT phosphorylation at S473 (AKT-pS473), was normal at 4 weeks of HFD (**Fig. S4E**). Shifting the diet from HFD to ND for 2 weeks restored HK2 expression and normal blood glucose levels (**Fig. 1C** and **Fig. S4D**), suggesting that loss of HK2 upon HFD is a transient physiological response. Furthermore, the finding that HK2 expression inversely correlated with hyperglycemia suggests that loss of adipose HK2 may be causal for loss of glucose homeostasis. Omental WAT (human vWAT) biopsies from obese non-diabetic and obese diabetic patients displayed reduced HK activity (**Fig. 1D** and **Table S2**), consistent with the previous observation that HK2 expression is decreased in adipose tissue of diabetic patients (*16*). These findings suggest that HK2 down-regulation in adipose tissue is a key event, possibly causal, in obesity-induced insulin insensitivity and hyperglycemia in mouse and human.

**Fig. 1.**
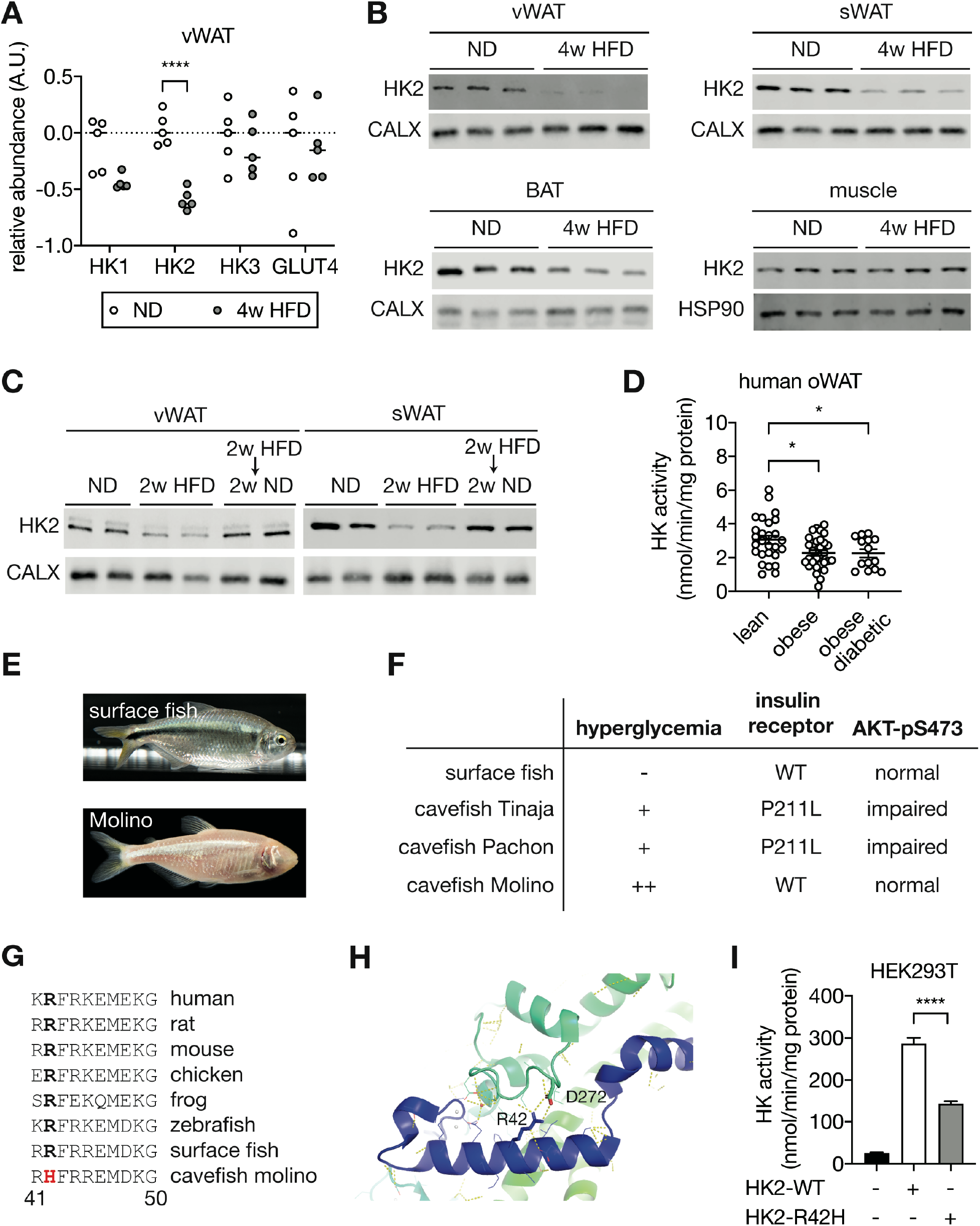
Loss of HK2 in HFD-fed mice, obese or/and diabetic humans, and cavefish Molino. (**A**) Relative protein levels of Hexokinases and GLUT4 in visceral white adipose tissue (vWAT) of normal diet (ND)- and 4-week high fat diet (HFD)-fed wild-type C57BL/6JRj mice. Multiple t test, ****p<0.0001. n=5 (ND) and 5 (HFD). (**B**)Immunoblot analyses of HK2 expression of vWAT, subcutaneous WAT (sWAT), brown adipose tissue (BAT), and skeletal muscle from ND- and 4-week HFD-fed mice. CALX and HSP90 serve as loading controls. n >10 (ND) and >10 (HFD). (**C**) Immunoblot analyses of HK2 expression in vWAT and sWAT from ND-, 2-week HFD-, and 2-week HFD + 2-week ND-fed mice. n=5 (ND), 5 (HFD), and 6 (HFD+ND). (**D**) HK activity of omental WAT (oWAT) from lean, obese, and obese diabetic patients. One-way ANOVA, *p<0.05. n=27 (lean), 30 (obese), and 14 (obese diabetic). (**E**) Surface fish and Mexican cavefish Molino. (**F**) Comparisons of phenotypes in surface fish, Pachón, Tinaja, and Molino. (**G**) Amino acid sequence alignment of the HK2-R42H mutation within vertebrates. (**H**) Structural analyses revealed the presence of a salt bridge between Arginine 42 (R42) and Aspartic acid 272 (D272) in the human HK2 (PDB: 2MTZ). (**I**) HK activity in lysates of HEK293T cells expressing control, HK2-WT, or HK2-R42H. Student’s t test, ****p<0.0001. N=4.

Mexican cavefish (*Astyanax mexicanus*), also known as blind fish, are hyperglycemic compared to surface fish from which they are descended (*17*) (**Fig. 1E-F**). Hyperglycemia, although a pathological condition in mouse and human, is a selected trait that allows cave-dwelling fish to survive in nutrient-limited conditions. Among three independently evolved cavefish isolates, Pachón and Tinaja cavefish (names refer to the caves from which the fish were isolated) contain a loss-of-function mutation in the insulin receptor, causing insulin resistance and hyperglycemia (*17*) (**Fig. 1F**). The third isolate, Molino cavefish, is the most hyperglycemic but contains a wild-type insulin receptor and displays normal insulin signaling (*17*). Since the phenotype of Molino fish is similar to that of HFD-fed mice, we hypothesized that this cavefish may be hyperglycemic due to a loss-of-function mutation in the *hk2* gene. DNA sequencing of the *hk2* gene of surface fish and the three cavefish variants revealed a mutation in the *hk2* gene uniquely in Molino. The homozygous, missense mutation in the coding region of the Molino *hk2* gene changed highly conserved arginine 42 (R42) to histidine (R42H) (**Fig. 1G** and **Fig. S5A**). Based on the published structure of HK2 (*18*), R42 forms a salt bridge with aspartic acid 272 (D272) to stabilize the conformation of HK2 (**Fig. 1H**), predicting that R42H destabilizes HK2 and is thus a loss-of-function mutation. To test this prediction, we over-expressed wild-type HK2 (HK-WT) and HK2 harboring the Molino mutation (HK2-R42H) in HEK293T cells which have low intrinsic hexokinase activity. HK2-R42H displayed ~50% lower hexokinase activity compared to HK2-WT, despite similar expression levels, suggesting that R42H is indeed a loss-of-function mutation (**Fig. 1I** and **Fig. S5B**). Thus, the R42H mutation may account for the hyperglycemia in Molino cavefish. In other words, R42H in Molino appears to be a genetically fixed version of what we observe in mice as a physiological down-regulation of HK2 in response to HFD. The findings in Molino provide orthogonal evidence that the down-regulation of HK2 in mice is physiologically relevant.

The above findings altogether suggest that adipose-specific loss of HK2 may be a cause of insulin insensitivity and hyperglycemia. To test this hypothesis, we first generated a stable HK2 knockdown pre-adipocyte 3T3-L1 cell line (**Fig. S6A-B**). HK2 knockdown pre-adipocytes differentiated normally to produce mature adipocytes (**Fig. S6C-D**). The glycolytic rate was lower in HK2 knockdown adipocytes compared to controls, as measured by extracellular acidification rate (ECAR) (**Fig. 2A**) and lactate production (**Fig. 2B**). Furthermore, although basal glucose uptake did not differ, insulin-stimulated glucose uptake was 50% lower in HK2 knockdown adipocytes (**Fig. 2C**). Thus, loss of HK2 causes insulin insensitivity in adipocytes *in vitro*.

**Fig. 2.**
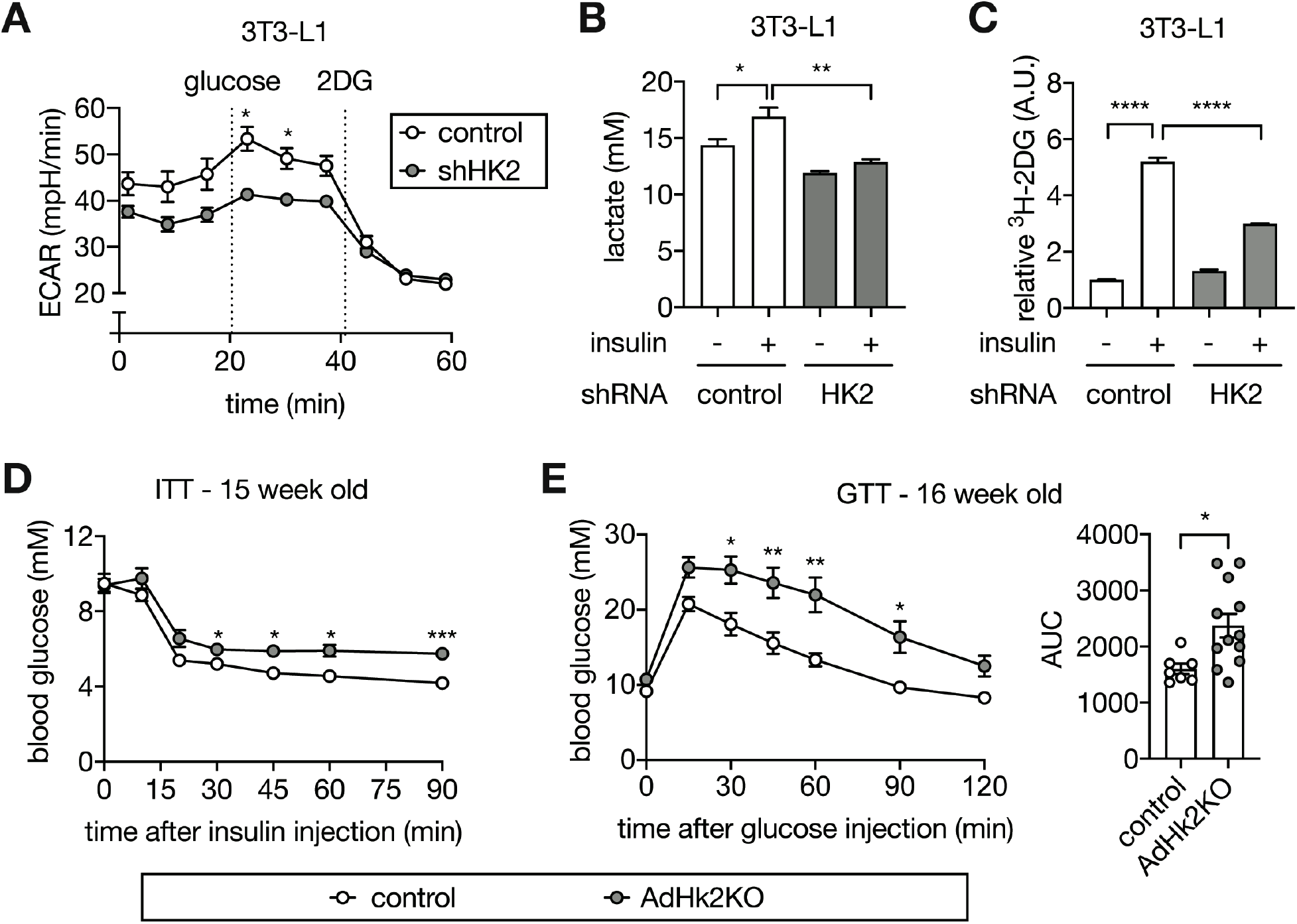
Loss of adipose HK2 causes glucose intolerance. (**A**) Extracellular acidification rate (ECAR) of control and HK2 knockdown 3T3-L1 adipocytes in response to glucose (10 mM) and 2-deoxyglucose (2DG, 50 mM). Two-way ANOVA. *p<0.05. N=8. (**B**) Lactate secreted into media by control and HK2 knockdown 3T3-L1 adipocytes treated with or without 100 nM insulin. One-way ANOVA. *p<0.05, **p<0.01. N=3. (**C**) 2DG uptake in control and HK2 knockdown 3T3-L1 adipocytes treated with or without 100 nM insulin. One-way ANOVA. ****p<0.0001. N=3. (**D**) Insulin tolerance test (ITT) on control and AdHk2KO mice. Mice were fasted for 6 hours and injected with insulin (0.5 U/kg body weight). Two-way ANOVA. *p<0.05, ***p<0.001. n=6 (control) and 10 (AdHk2KO). (**E**) Gucose tolerance test (GTT) on control and AdHk2KO mice. Mice were fasted for 6 hours and injected with glucose (2 g/kg body weight). Two-way ANOVA for glucose curves and Student’s t test for AUC. *p<0.05, **p<0.01. n=7 (control) and 12 (AdHk2KO).

To examine further the role of adipose HK2 in glucose homeostasis, and in particular the causality of HK2 loss in hyperglycemia, we generated an adipose-specific *Hk2* knockout (AdHk2KO) mouse. In AdHk2KO mice, HK2 expression was decreased in vWAT, sWAT, and BAT but unchanged in skeletal muscle (**Fig. S7** and **S8**), similar to HFD-fed mice (**Fig. 1B**). AdHk2KO mice displayed slightly less fat mass and slightly more lean mass than controls, but no difference in overall body weight (**Fig. S9A-C**). No significant difference was observed in organ weight, except for a small decrease in vWAT and a small increase in liver weight (**Fig. S9D-E).** Importantly, AdHk2KO mice were insulin insensitive and severely glucose intolerant (**Fig. 2D** and **2E**). These findings indicate that loss of HK2 specifically in adipose tissue causes insulin insensitivity and hyperglycemia.

Adipose tissue accounts for only <5% of glucose disposal (*3, 4*). Thus, the severe glucose intolerance observed in AdHk2KO mice (**Fig. 2E**) cannot be explained solely by impaired glucose uptake into adipose tissue. Adiponectin and leptin, two well-known adipokines controlling systemic glucose homeostasis (*19*), were unchanged in AdHk2KO mice compared to controls (**Fig. S10A-E**). Another possible explanation for the observed severity of glucose intolerance could be impaired insulin secretion. However, plasma insulin levels were similar in AdHk2KO and control mice (**Fig. S11A-B**). Moreover, insulin signaling was not affected in adipose tissue, skeletal muscle, and liver of fasted and re-fed AdHk2KO mice (**Fig. S7, S8**, and **S12A**). These findings indicate that hyperglycemia in AdHk2KO mice is not due to defective adipokine expression or insulin signaling.

Another potential explanation for the severity of glucose intolerance in AdHk2KO mice is de-repressed hepatic glucose production. Adipose tissue is known to impinge negatively on glucose production in liver (*6, 9, 11, 20*), the main glucose producing organ. To examine whether AdHk2KO mice display enhanced hepatic glucose production, we first performed a pyruvate tolerance test (PTT). AdHk2KO mice displayed significantly higher glucose production compared to controls (**Fig. 3A**). We next examined expression of gluconeogenic genes (*G6pc* and *Pepck*) in liver. Consistent with the observed increase in glucose production, gluconeogenic genes were upregulated in AdHk2KO liver (**Fig. 3B**). These findings suggest that enhanced hepatic gluconeogenesis accounts for the severity of glucose intolerance in AdHk2KO mice. Thus, loss of adipose HK2 non-cell-autonomously promotes hyperglycemia by enhancing hepatic gluconeogenesis.

**Fig. 3.**
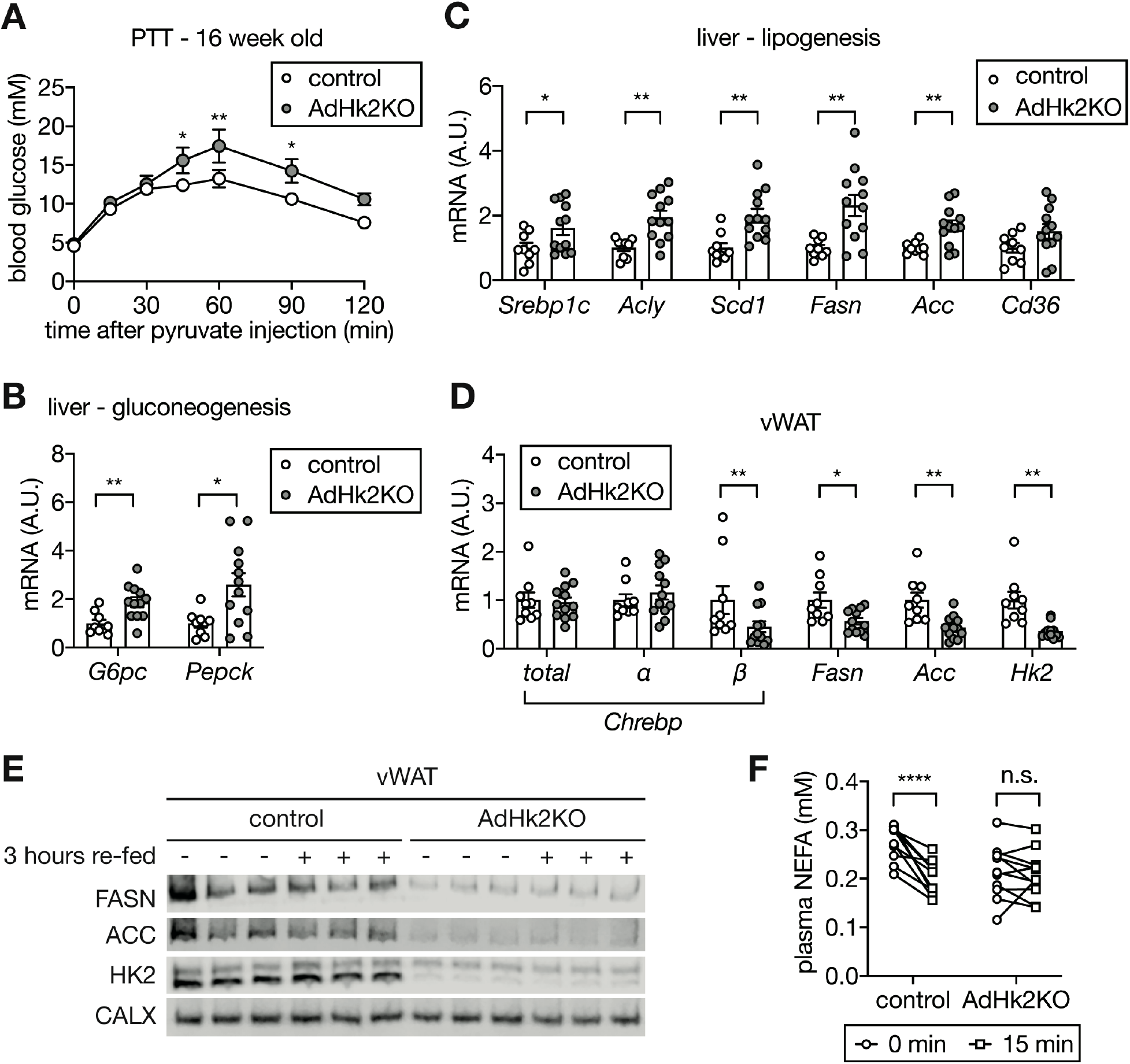
Loss of adipose HK2 promotes hepatic gluconeogenesis by inhibiting ChREBP and lipogenesis in adipose tissue. (**A**) Pyruvate tolerance test (PTT) on control and AdHk2KO mice. Mice were fasted for 15 hours and injected with pyruvate (2 g/kg body weight). Two-way ANOVA, *p<0.05, **p<0.01. n=5 (control) and 6 (AdHk2KO). (**B**) mRNA levels of gluconeogenic genes in liver of control and AdHk2KO mice. Multiple t test, *p<0.05, **p<0.01. n=9 (control) and 12 (AdHk2KO). (**C**) mRNA levels of lipogenic genes in liver from control and AdHk2KO mice. Multiple t test, *p<0.05, **p<0.01. n=9 (control) and 12 (AdHk2KO). (**D**) mRNA levels of lipogenic genes in vWAT from control and AdHk2KO mice. Multiple t test, *p<0.05, **p<0.01. n=9 (control) and 12 (AdHk2KO). (**E**) Immunoblots analyses of lipogenic enzymes in vWAT of control and Adk2KO mice. n=6 (fasted control), 6 (re-fed control), 6 (fasted AdHk2KO), and 6 (re-fed AdHk2KO). (**F**) Plasma non-esterified fatty acid (NEFA) levels in control and AdHk2KO mice. Mice were fasted for 6 hours and injected with glucose (2g/kg body weight). Two-way ANOVA, ****p<0.0001. n=10 (control) and 11 (AdHk2KO).

We observed enhanced gluconeogenic capacity in AdHk2KO mice despite normal systemic insulin signaling, including in liver. To investigate this apparent contradiction, we examined lipogenic gene expression, another readout for hepatic insulin action. Sterol regulatory element-binding protein 1c (SREBP1c) is a transcription factor that activates lipogenic genes and thereby promotes *de novo* lipogenesis in the liver (*21*). Expression of SREBP1c and its target fatty acid synthesis genes (*Acly*, *Scd1*, *Fasn*, and *Acc*) were increased in liver of AdHk2KO mice (**Fig. 3C**). FASN and ACC protein levels were also increased in liver of AdHk2KO mice (**Fig. S12A**). These data further suggest that loss of adipose HK2 promotes hepatic lipogenesis. The increased hepatic gluconeogenic and lipogenic capacity observed in AdHk2KO mice mimics a condition in diabetic patients known as selective insulin resistance (*22*) (see below). Thus, loss of adipose HK2 causes selective insulin resistance and thereby contributes to the pathogenesis of type 2 diabetes.

We note that hepatic triglyceride (TG) levels were unchanged in AdHk2KO mice despite increased expression of lipogenic genes and proteins (**Fig. 3C** and **Fig. S12A-B**). Most likely, *de novo*-synthesized TG in AdHk2KO liver is delivered to adipose tissue for storage. This is consistent with our observation that adipose tissue mass is largely unaffected in AdHk2KO mice.

How does loss of adipose HK2 increase hepatic gluconeogenesis? Adipose lipogenesis non-cell-autonomously inhibits hepatic glucose production (*20, 23, 24*). More specifically, adipose-specific knockout of the transcription factor carbohydrate-responsive element binding protein (ChREBP) decreases adipose lipogenesis and thereby increases hepatic gluconeogenesis (*20, 25*). Furthermore, HFD inhibits adipose ChREBP and lipogenesis via an unknown mechanism proposed to involve glucose or a glucose metabolite (*26*). G6P, a glucose metabolite and product of HK2, promotes ChREBP activity in a rat insulinoma cell line (*27*). Thus, we hypothesized that HFD-induced loss of HK2 and a concomitant decrease in G6P leads to loss of both ChREBP activity and lipogenesis in adipose tissue. To test this hypothesis, we examined *Chrebp* expression in adipose tissue of AdHk2KO and control mice. ChREBP has two isoforms. Constitutively expressed ChREBP*α* promotes transcriptional activation of ChREBP*β* which then activates lipogenic genes (*26*). Expression of *Chrebpβ,* but not *Chrebpα*, was significantly decreased in vWAT, sWAT, and BAT of AdHk2KO mice (**Fig. 3D** and **S13A-B**). Consistent with reduced *Chrebpβ* expression, lipogenic genes and proteins were down-regulated in adipose tissue of AdHk2KO mice (**Fig. 3D-E** and **S13A-B**). These findings suggest that loss of adipose HK2 increases hepatic gluconeogenesis via loss of ChREBP activity and lipogenesis in adipose tissue.

Consistent with the role of adipose lipogenesis in inhibiting hepatic gluconeogenesis, adipose lipolysis and consequential release of non-esterified fatty acid (NEFA) enhances glucose production in liver (*23*). Furthermore, in addition to activating lipogenic genes, ChREBP inhibits NEFA release (*20*). We examined the effect of HK2 loss on NEFA release. Glucose administration increased insulin levels in fasted AdHk2KO and control mice in a similar manner (**Fig. S11B**). However, this physiological increase in insulin levels inhibited NEFA release in control mice but not in AdHk2KO mice (**Fig. 3F**). These findings combined with previous observations (*11, 23, 24*) suggest that loss of HK2 promotes release of NEFA which in turn de-represses hepatic gluconeogenesis. Thus, loss of adipose HK2 both prevents storage and promotes release of fatty acid that activates gluconeogenesis in the liver.

Our findings suggest that HFD causes loss of adipose HK2 and thereby triggers hyperglycemia. How does HFD down-regulate adipose HK2? To answer this question, we first examined *Hk2* mRNA expression in HFD- and ND-fed mice. HFD-fed mice exhibited decreased *Hk2* mRNA in sWAT but not in vWAT or BAT (**Fig. 4A**). This suggests that HK2 synthesis is down-regulated at the transcriptional level in sWAT, and at a post-transcriptional level in vWAT and BAT. To investigate the post-transcriptional mechanism, we performed polysome profiling on vWAT from HFD- and ND-fed mice. In the presence of cycloheximide to maintain ribosome-mRNA complexes, vWAT lysates from HFD-fed mice contained more polysomes and fewer monosomes (80S ribosome) relative to ND-fed mice (**Fig. 4B**). An increase in polysomes can be due to increased translation initiation or decreased elongation. To distinguish these two possibilities, we performed polysome profiling in the absence of cycloheximide, a condition which allows actively translating ribosomes to run off mRNA. Although there were fewer overall polysomes in the absence of cycloheximide, we consistently observed more polysomes in vWAT from HFD-fed mice (**Fig. 4C**), suggesting that HFD inhibits or slows translation elongation of a subset of mRNA species. Shifting the diet from HFD to ND for 2 weeks restored elongation (**Fig. 4D**), as described above for glycemia (**Fig. S4D**). To determine whether HFD affects elongation of *Hk2* mRNA translation in particular, we measured *Hk2* mRNA in monosome, light polysome, and heavy polysome fractions. *Hk2* mRNA from HFD vWAT was enriched in heavy polysomes, whereas it was enriched in light polysomes in ND vWAT (**Fig. 4E** and **Fig. S14A**). To further examine reduced HK2 protein synthesis, we measured synthesis of HK2 in vWAT of HFD-fed mice. vWAT explants from HFD- or ND-fed mice were treated with L-azidohomoalanine (AHA), a methionine analog, and AHA-containing polypeptides were purified and quantified by mass spectrometry. The amount of AHA-containing HK2 polypeptides was significantly decreased in vWAT explants from HFD-fed mice (**Fig. 4F**). Thus, HFD down-regulates HK2 in vWAT by inhibiting *Hk2* mRNA translation.

**Fig. 4.**
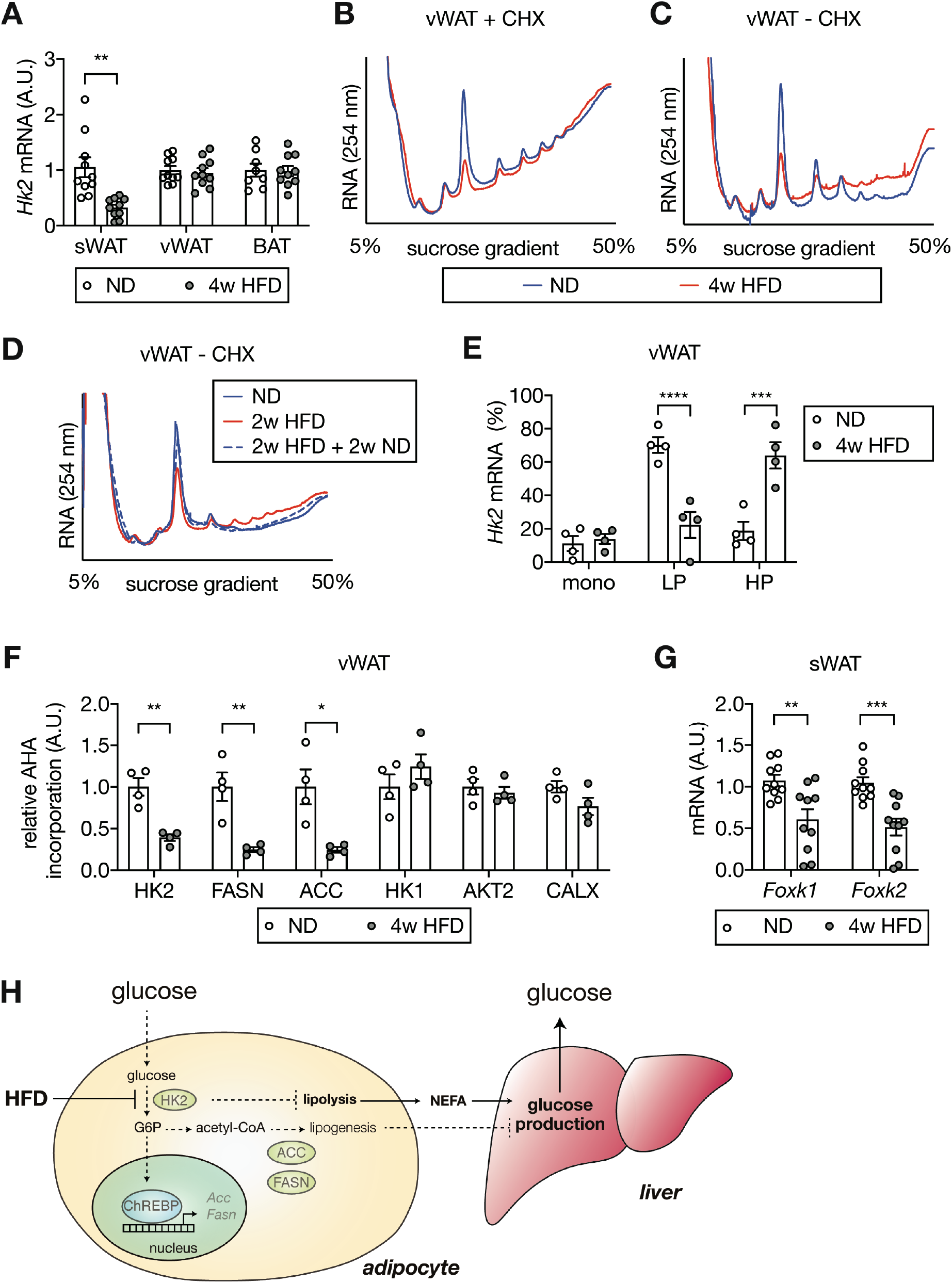
Mechanism of HFD-induced HK2 down-regulation in adipose tissue. (**A**) *Hk2* mRNA levels in sWAT, vWAT, and BAT from ND- and 4-week HFD-fed mice. Student’s t test, **p<0.01. n=10 (sWAT from ND and HFD), 10 (vWAT from ND and HFD), 8 (BAT from ND), 10 (BAT from HFD). (**B**) Representative polysome profiles of vWAT from ND- and 4-week HFD-fed mice. Tissue lysates were prepared in the presence of cycloheximide (CHX) to prevent ribosome run-off. n=6 (ND) and 6 (HFD). (**C**) Representative run-off polysome profiles of vWAT from ND- and 4-week HFD-fed mice. Tissue lysates were prepared in the absence of CHX to allow run-off. n=7 (ND) and 7 (HFD). (**D**) Representative run-off polysome profiles of vWAT from ND-, 2-week HFD-, 2-week HFD + 2-week ND-fed mice. Tissue lysates were prepared in the absence of CHX. n=5 (ND) and 5 (HFD). (**E**) *Hk2* mRNA abundance in monosome (mono), light polysome (LP), and heavy polysome (HP) fractions obtained in Fig. 4C. Two-way ANOVA, ***p<0.001, ****p<0.0001. n=4 (ND) and 4 (HFD). (**F**) Nascent polypeptides. vWATs isolated from ND- 4-week HFD-fed mice were labeled with AHA, and AHA-containing peptides were measured by mass spectrometer. *p<0.05, **p<0.01. n=4 (ND) and 4 (HFD). (**G**) *Foxk1* and *Foxk2* mRNA levels in sWAT from ND- or 4-week HFD-fed mice. Multiple t test, **p<0.01, ***p<0.001. n=10 (ND) and 10 (HFD). (**H**) Diet-induced loss of adipose HK2 triggers hyperglycemia via reduced glucose clearance in adipocytes (left) and de-repressed hepatic gluconeogenesis (right).

It was recently reported that the forkhead transcription factors FOXK1 and FOXK2 promote *Hk2* transcription in adipose tissue(*28*). We investigated whether the transcriptional down-regulation of *Hk2* mRNA in sWAT is due to loss of FOXK1 and FOXK2. *Foxk1* and *Foxk2* expression was significantly decreased in sWAT, but not in vWAT or BAT, of HFD-fed mice (**Fig. 4G** and **S14B-C**). Thus, HFD appears to down-regulate HK2 in sWAT by inhibiting FOXK1/2 and, thereby, expression of the *Hk2* gene.

How obesity causes hyperglycemia is a long-standing question. In this study, we show that a high fat diet causes hyperglycemia by inducing loss of HK2 specifically in adipose tissue. Loss of adipose HK2 leads to hyperglycemia in two ways (**Fig. 4H**). First, HK2 loss results in reduced glucose uptake by adipose tissue due to the inability of adipocytes to trap non-phosphorylated glucose. Second, loss of HK2 in adipocytes de-represses glucose production in liver.

In liver, insulin normally down-regulates gluconeogenesis to decrease blood glucose and up-regulates lipogenesis to increase energy storage. Paradoxically, in type 2 diabetes, insulin fails to inhibit gluconeogenesis but stimulates lipogenesis, hence the liver is selectively insulin resistant (*22*). Furthermore, previous findings suggest that selective resistance occurs despite intact hepatic insulin signaling (*24*). Consistent with these previous findings, we observed that loss of adipose HK2 causes selective insulin resistance without affecting systemic insulin signaling. Thus, we propose that loss of adipose HK2 is a mechanism for selective insulin resistance and ultimately diabetes.

We found an HK2-R42H mutation in naturally hyperglycemic Mexican cavefish. To date, no human monogenic disease has been reported associated with mutations in the *HK2* gene. However, 12 R42W and 4 R42Q alleles are reported in the genomAD database(*29*). Further studies are required to determine whether these alleles are loss-of-function and associated with diabetes.

## Acknowledgements

We thank Stefan Offermanns (MPI-HLR, Germany), Didier Trono (EPFL, Switzerland), Robert Weinberg (MIT, USA), Christine Riggenbach (St. Claraspital), Christoph Handschin (Biozentrum), the Imaging Core Facility (Biozentrum), and the Proteomics Core Facility (Biozentrum) for providing reagents and technical support.

## Author contributions

MS and MNH conceived the project and designed the experiments with contributions from TM and NR. MS, SS, ICF, DW, ND, AT, DR, NR performed experiments and analyzed data. NH provided reagents and mice. BKW, ACM, and RP contributed to the collection of human adipose tissue biopsies. MS and MNH wrote the manuscript.

## Supplementary Materials

Materials and Methods

Figs S1-S14

Tables S1-S2

References (30-36)

## Materials and Methods

### Mouse

Wild-type C57BL/6JRj mice were purchased from JANVIER LABS. Mice carrying *Hk2* with exons 4-10 flanked by loxP sites (*Hk2^fl/fl^*) were as described (*30*). *Adipoq-CreER^T2^* mice were provided by Prof. Stefan Offermanns (MPI-HLR, Germany) (*31*). *Hk2^fl/fl^* mice were crossed with *Adipoq-CreER^T2^* mice, and resulting Cre positive *Hk2^fl/+^* mice were crossed with *Hk2^fl/fl^* mice to generate adipocyte-specific *Hk2* knockout (*Adipoq-CreER^T2^* positive *Hk2^fl/fl^*) mice (AdHk2KO). *Hk2* knockout was induced by *i.p. injection of* 1 mg/mouse tamoxifen (Sigma-Aldrich) in corn oil for 7 days. Littermate *Cre* negative animals were used as a control. Control mice were also treated with tamoxifen.

Mice were housed at 22 °C in a conventional facility with a 12 hour light/dark cycle with unlimited access to water, and normal diet (ND) or high fat diet (HFD: 60% kcal % fat NAFAG 2127, KLIBA). Only male mice between 6 and 17 weeks of age were used for experiments.

### Plasmids

The R42H mutation was introduced into the pLenti-CMV-ratHK2 by PCR using the oligos 5’atttctaggcACttccggaaggagatggagaaag3’ and 5’cttccggaaGTgcctagaaatctccagaagggtc3’. The desired sequence change was confirmed.

### Cell culture

HEK293T cells were obtained from ATCC and cultured in M1 (DMEM supplemented with 4 mM glutamine, 1mM sodium pyruvate, 1x penicillin and streptomycin, and 10% FBS). 3T3-L1 cells were obtained from ATCC and cultured and differentiated as previously described(*32*). In brief, 3T3-L1 preadipocyte cells were maintained in M1 medium at 37 °C incubator with 5% CO_2_. For differentiation, cells were maintained in M1 medium for 2 days after reaching confluence. The cells were then transferred to M2 medium (M1 medium supplemented with 1.5 μg/mL insulin, 0.5 mM IBMX, 1 μM dexamethasone, and 2 μM rosiglitazone), defined as day 0 post-differentiation. After 2 days, the cells were transferred to M3 medium (M1 with 1.5 μg/mL insulin). At day 4 post-differentiation, cells were transferred back to M2 medium. From day 6 post-differentiation, cells were maintained in M3 with medium change every two days.

For Hk2 knockdown, MISSION shRNA (TRCN0000280118) or control pLKO plasmid were purchased from Merck and co-transfected with psPAX2 (a gift from Didier Trono: Addgene plasmid # 12260) and pCMV-VSV-G (*33*) (a gift from Robert Weinberg: Addgene plasmid # 8454) into HEK293T cells. Supernatants containing lentivirus were collected one day after transfection, and used to infect undifferentiated 3T3-L1 cells. Transduced cells were selected by puromycin. For all experiments, 8-14 days post-differentiated cells were used.

### Human biopsies

Omental white adipose tissue (oWAT) biopsies were obtained from lean subjects with normal fasting glucose level and body mass index (BMI) < 27 kg/m^2^, from obese non-diabetic subjects with BMI > 35 kg/m^2^ HbA1c < 6.0%, and from obese diabetic sujects with BMI > 35 kg/m^2^ HbA1c > 6.1% (**Table S2**). All subjects gave informed consent before the surgical procedure. Patients were fasted overnight and underwent general anesthesia. All oWAT specimens were obtained between 8:30 and 12:00am, snap-frozen in liquid nitrogen, and stored at −80 °C for subsequent use.

### Proteomics

Tissues were pulverized and homogenized in lysis buffer containing 100 mM Tris-HCl pH7.5, 2 mM EDTA, 2 mM EGTA, 150 mM NaCl, 1% Triton X-100, cOmplete inhibitor cocktail (Roche) and PhosSTOP (Roche). Proteins were precipitated by trichloroacetic acid (Sigma) and the resulting protein pellets were washed with cold acetone. 25 μg of peptides were labeled with isobaric tandem mass tags (TMT 10-plex, Thermo Fisher Scientific) as described previously (*34*). Shortly, peptides were resuspended in 20 μl labeling buffer (2 M urea, 0.2 M HEPES, pH 8.3) and 5 μL of each TMT reagent were added to the individual peptide samples followed by a 1 h incubation at 25 °C. To control for ratio distortion during quantification, a peptide calibration mixture consisting of six digested standard proteins mixed in different amounts was added to each sample before TMT labeling. To quench the labelling reaction, 1.5 μL aqueous 1.5 M hydroxylamine solution was added and samples were incubated for another 10 min at 25 °C followed by pooling of all samples. The pH of the sample pool was increased to 11.9 by adding 1 M phosphate buffer (pH 12) and incubated for 20 min at 25 °C to remove TMT labels linked to peptide hydroxyl groups. Subsequently, the reaction was stopped by adding 2 M hydrochloric acid and until a pH < 2 was reached. Finally, peptide samples were further acidified using 5% TFA, desalted using Sep-Pak Vac 1cc (50 mg) C18 cartridges (Waters) according to the manufacturer’s instructions and dried under vacuum.

TMT-labeled peptides were fractionated by high-pH reversed phase separation using a XBridge Peptide BEH C18 column (3,5 μm, 130 Å, 1 mm × 150 mm, Waters) on an Agilent 1260 Infinity HPLC system. Peptides were loaded on column in buffer A (20 mM ammonium formate in water, pH 10) and eluted using a two-step linear gradient from 2% to 10% in 5 minutes and then to 50% buffer B (20 mM ammonium formate in 90% acetonitrile, pH 10) over 55 min at a flow rate of 42 μl/min. Elution of peptides was monitored with a UV detector (215 nm, 254 nm) and a total of 36 fractions were collected, pooled into 12 fractions using a post-concatenation strategy as previously described(*35*) and dried under vacuum.

Dried peptides were resuspended in 20 μl of 0.1% aqueous formic acid and subjected to LC–MS/MS analysis using a Q Exactive HF Mass Spectrometer fitted with an EASY-nLC 1000 (both Thermo Fisher Scientific) and a custom-made column heater set to 60 °C. Peptides were resolved using a RP-HPLC column (75 μm × 30 cm) packed in-house with C18 resin (ReproSil-Pur C18–AQ, 1.9 μm resin; Dr. Maisch GmbH) at a flow rate of 0.2 μL/min. The following gradient was used for peptide separation: from 5% B to 15% B over 10 min to 30% B over 60 min to 45 % B over 20 min to 95% B over 2 min followed by 18 min at 95% B. Buffer A was 0.1% formic acid in water and buffer B was 80% acetonitrile, 0.1% formic acid in water.

The mass spectrometer was operated in DDA mode with a total cycle time of approximately 1 sec. Each MS1 scan was followed by high-collision-dissociation (HCD) of the 10 most abundant precursor ions with dynamic exclusion set to 30 sec. For MS1, 3e6 ions were accumulated in the Orbitrap over a maximum time of 100 ms and scanned at a resolution of 120,000 FWHM (at 200 m/z). MS2 scans were acquired at a target setting of 1e5 ions, accumulation time of 100 ms and a resolution of 30,000 FWHM (at 200 m/z). Singly charged ions and ions with unassigned charge state were excluded from triggering MS2 events. The normalized collision energy was set to 35%, the mass isolation window was set to 1.1 m/z and one microscan was acquired for each spectrum.

The acquired raw-files were converted to the mascot generic file (mgf) format using the msconvert tool (part of ProteoWizard, version 3.0.4624 (2013-6-3)). Using the MASCOT algorithm (Matrix Science, Version 2.4.1), the mgf files were searched against a decoy database containing normal and reverse sequences of the predicted SwissProt entries of Mus musculus (www.ebi.ac.uk, release date 2014/11/24), the six calibration mix proteins(*34*) and commonly observed contaminants (in total 50214 sequences for Mus musculus) generated using the SequenceReverser tool from the MaxQuant software (Version 1.0.13.13). The precursor ion tolerance was set to 10 ppm and fragment ion tolerance was set to 0.02 Da. The search criteria were set as follows: full tryptic specificity was required (cleavage after lysine or arginine residues unless followed by proline), 3 missed cleavages were allowed, carbamidomethylation (C) and TMT6plex (K and peptide N-terminus) were set as fixed modification and oxidation (M) as a variable modification. Next, the database search results were imported into the Scaffold Q+ software (version 4.3.2, Proteome Software Inc., Portland, OR) and the protein false identification rate was set to 1% based on the number of decoy hits. Proteins that contained similar peptides and could not be differentiated based on MS/MS analysis alone were grouped to satisfy the principles of parsimony. Proteins sharing significant peptide evidence were grouped into clusters. Acquired reporter ion intensities in the experiments were employed for automated quantification and statically analysis using a modified version of our in-house developed SafeQuant R script(*34*). This analysis included adjustment of reporter ion intensities, global data normalization by equalizing the total reporter ion intensity across all channels, summation of reporter ion intensities per protein and channel, calculation of protein abundance ratios and testing for differential abundance using empirical Bayes moderated t-statistics. Finally, the calculated p-values were corrected for multiple testing using the Benjamini−Hochberg method (q-value). Deregulated proteins were selected by log2(fold change) > 0.6 or log2(fold change) < −0.6, q-value < 0.01.

### Immunoblots

Adipose tissue, liver, and skeletal muscle were homogenized in lysis buffer containing 100 mM Tris-HCl pH7.5, 2 mM EDTA, 2 mM EGTA, 150 mM NaCl, 1% Triton X-100, cOmplete inhibitor cocktail (Roche) and PhosSTOP (Roche). Protein concentration was determined by the Bradford assay, and equal amounts of protein were separated by SDS-PAGE, and transferred onto nitrocellulose membranes (GE Healthcare). Antibodies used in this study were as follows: AKT (Cat#4685 or Cat#2920), AKT-pS473 (Cat#4060), AKT-pT308 (Cat#13038), PRAS40-pT246 (Cat#2997), PRAS40 (Cat#2691), HK2 (Cat#2867), HK1 (Cat#2024), IR-pY1146 (Cat#3021), IR (Cat#3025), S6K-pT389 (Cat#9234), S6K (Cat#2708), S6-pS240/244 (Cat#5364), S6 (Cat#2217), GSK3-pS21 (Cat# 5676), GSK3 (Cat#9331), FASN (Cat#3189), ACC (Cat#3662), HA tag (Cat#3724) from Cell Signaling, GLUT4 (Cat# NBP2-22214, NOVUS), HSP90, (Cat#sc-13119, Santa Cruz Biotechnology), CALNEXIN (Cat#ADI-SPA-860-F, Enzo Life Sciences) and ACTIN (Cat#MAB1501, Millipore).

### RNA isolation and quantitative RT-PCR

Total RNA was isolated with TRIzol reagent (Sigma-Aldrich) and RNeasy kit (Qiagen). RNA was reverse-transcribed to cDNA using iScript cDNA synthesis kit (BIO-RAD). Semiquantitative real-time PCR analysis was performed using fast SYBR green (Applied Biosystems). Relative expression levels were determined by normalizing each CT values to *Tbp* using the ΔΔCT method. The sequence for the primers used in this study was as follows. Fasn-fw: 5’GCTGCGGAAACTTCAGGAAAT3’, Fasn-rv: 5’AGAGACGTGTCACTCCTGGACTT3’, Acc-fw: 5’AAGGCTATGTGAAGGATG3’, Acc-rv: 5’CTGTCTGAAGAGGTTAGG3’, Cd36-fw: 5’ TGGCCTTACTTGGGATTGG3’, Cd36-rv: 5’ CCAGTGTATATGTAGGCTCATCCA 3’, G6pc-fw: 5’CCATGCAAAGGACTAGGAACAA3’, G6pc-rv: 5’ TACCAGGGCCGATGTCAAC 3’, Pepck-fw: 5’ CCACAGCTGGTGCAGAACA3’, Pepck-rv: 5’ GAAGGGTCGATGGCAAA 3’, Chrebp-fw: 5’CACTCAGGGAATACACGCCTAC3’, Chrebp-rv: 5’ATCTTGGTCTTAGGGTCTTCAGG3’, Chrebp-alpha-fw: 5’CGACACTCACCCACCTCTTC3’, Chrebp-alpha-rv: 5’TTGTTCAGCCGGATCTTGTC3’, Chrebp-beta-fw: 5’TCTGCAGATCGCGTGGAG3’, Chrebp-beta-rv: 5’CTTGTCCCGGCATAGCAAC3’, Hk2-fw: 5’ ACGGAGCTCAACCAAAACCA3’, Hk2-rv: 5’TCCGGAACCGCCTAGAAATC3’, Foxk1-fw: 5’ GGCTGTCACTCAGAATGGAA3’, Foxk1-rv: 5’GAGGCAGATGTGGTAGTGGAG3’, Foxk2-fw: 5’ CCACGGGAACTATCAGTGCT3’, Foxk2-rv: 5’ GTCATCCTTTGGGCTGTCTC3’, Lep-fw: 5’ TCACACACGCAGTCGGTATC, Lep-rv: 5’ACTCAGAATGGGGTGAAGCC3’, Adipoq-fw: 5’TGACGACACCAAAAGGGCTC3’, Adipoq-rv: 5’ ACGTCATCTTCGGCATGACT3’, aP2-fw: 5’ TCGGTTCCTGAGGATACAAGAT3’, aP2-rv: 5’TTTGATGACTGTGGGATTGAAG3’, Tbp-fw: 5’TGCTGTTGGTGATTGT3’, Tbp-rv:5’ CTTGTGTGGGAAAGAT3’.

### Hexokinase assay

Measurements were performed with a hexokinase assay kit (abcam) following manufacturer’s instructions.

### Seahorse analyses

Measurements were performed with an XF96 Extracellular Flux Analyzer (Seahorse Bioscience of Agilent) following manufacturer’s instructions.

### Lactate measurement

Differentiated adipocytes were starved serum overnight and stimulated with 100 nM insulin for 2 hours. Extracellular lactate was measured with the Lactate Pro 2 analyzer (Axonlab).

### ^3^H-2DG uptake assay

Differentiated adipocytes were starved for serum for 5 hours and then incubated in Krebs Ringer Phosphate Hepes (KRPH) buffer with or without 100 nM insulin for 20 min. Cells were incubated with 50 μM 2DG containing 0.25 μCi ^3^H-2-deoxyglucose (2DG, Perkin Elmer) for 5 min and washed three times with cold PBS. Cells were lysed in lysis buffer and cleared by centrifugation at 14,000 g for 10 min. Incorporated ^3^H-2DG was measured with a scintillation counter.

### Insulin tolerance test, glucose tolerance test, pyruvate tolerance test

For the insulin and glucose tolerance tests, mice were fasted for 6 hours and insulin Humalog (Lilly, i.p. 0.75 or 0.5 U/kg body weight) or glucose (2 g/kg body weight) was given, respectively. For the pyruvate tolerance test, mice were fasted for 15 hours and pyruvate (2 g/kg body weight) was administered. Blood glucose was measured with a blood glucose meter (Accu-Check).

### Polysome profile analyses

Tissues were homogenized in polysome lysis buffer (10 mM Tris-HCl, pH7.5, 140 mM KCl, 5 mM MgCl_2_, 1mM DTT, 0.5% Triton x-100, 0.05% sodium deoxycholate, 40 u/mL RNase inhibitor) in the presence or absence of 100 μg/mL cycloheximide and the lysates were centrifuged twice at 14,000 g for 10 min. The cleared lysates were loaded onto 5-50% sucrose gradients (10 mM Tris-HCl, pH7.5, 140 mM KCl, 5 mM MgCl_2_, 5% or 50% sucrose) and centrifuged with TH-641 (Thermo Scientific) at 36,000 rpm for 2 hours at 4 °C. Fractions were collected by a Gradient Station (BIOCOMP INSTRUMENTS) with continuous recording of optical density (OD) at 254 nm. RNA from each fraction was isolated with TRIzol and qRT-PCR were performed as described(*36*).

### AHA-incorporation

vWATs from ND- or HFD-fed mice were immediately incubated in low glucose DMEM containing 50 μM azidohomoalanine (AHA) for 30. The tissues were powdered, lysed by bio-rupture and clarified by centrifugation at 15,000 g for 15 min twice. The 100 μg proteins were used for CLICK reaction following manufacturer’s instructions. The biotinylated proteins were pulled-down using streptavidin magnetic beads for 2 hours at 4 °C. The pulled-down proteins were digested with trypsin. The digested peptides were acidified using 5 % TFA and desalted using C18 columns. The eluted peptides were dried and analyzed by mass spectrometry.

### Insulin ELISA

Plasma insulin levels were measured by ultrasensitive mouse insulin ELISA kit (Crystal Chem) according to the manufacturer’s instructions.

### Leptin ELISA

Plasma Leptin levels were measured by mouse mouse Leptin ELISA kit (Crystal Chem) according to the manufacturer’s instructions.

### Adiponectin ELISA

Plasma Adiponectin levels were measured by mouse mouse Adiponectin ELISA kit (Crystal Chem) according to the manufacturer’s instructions.

### Hepatic triglyceride measurement

Hepatic triglyceride levels were measured using a triglyceride assay kit (Abcam) according to the manufacturer’s instructions.

### Body composition measurement

Body composition was measured by nuclear magnetic resonance imaging (Echo Medical Systems).

### Study Approval

All animal experiments were performed in accordance with federal guidelines for animal experimentation and were approved by the Kantonales Veterinäramt of the Kanton Basel-Stadt. For human biopsies, the study protocol was approved by the Ethikkomission Nordwest- und Zentralschweiz (EKNZ).

### Statistics

Sample size was chosen according to our previous studies and published reports in which similar experimental procedures were described. The investigators were not blinded to the treatment groups. All data are shown as the mean ± SEM. Sample numbers are indicated in each figure legend. For mouse experiments, *n* represents the number of animals, and for cell culture experiments, *N* indicates the number of independent experiments. To determine the statistical significance between 2 groups, an unpaired two-tailed Student’s t test was performed. For more than 3 groups, one-way ANOVA was performed. For ITT, GTT, PTT, weigh curve data, two-way ANOVA was performed. All statistical analysis was performed using GraphPad Prism 8 (GraphPad Software). A *p* value of less than 0.05 was considered statistically significant.

## Supplemental Figures

**Fig. S1.**
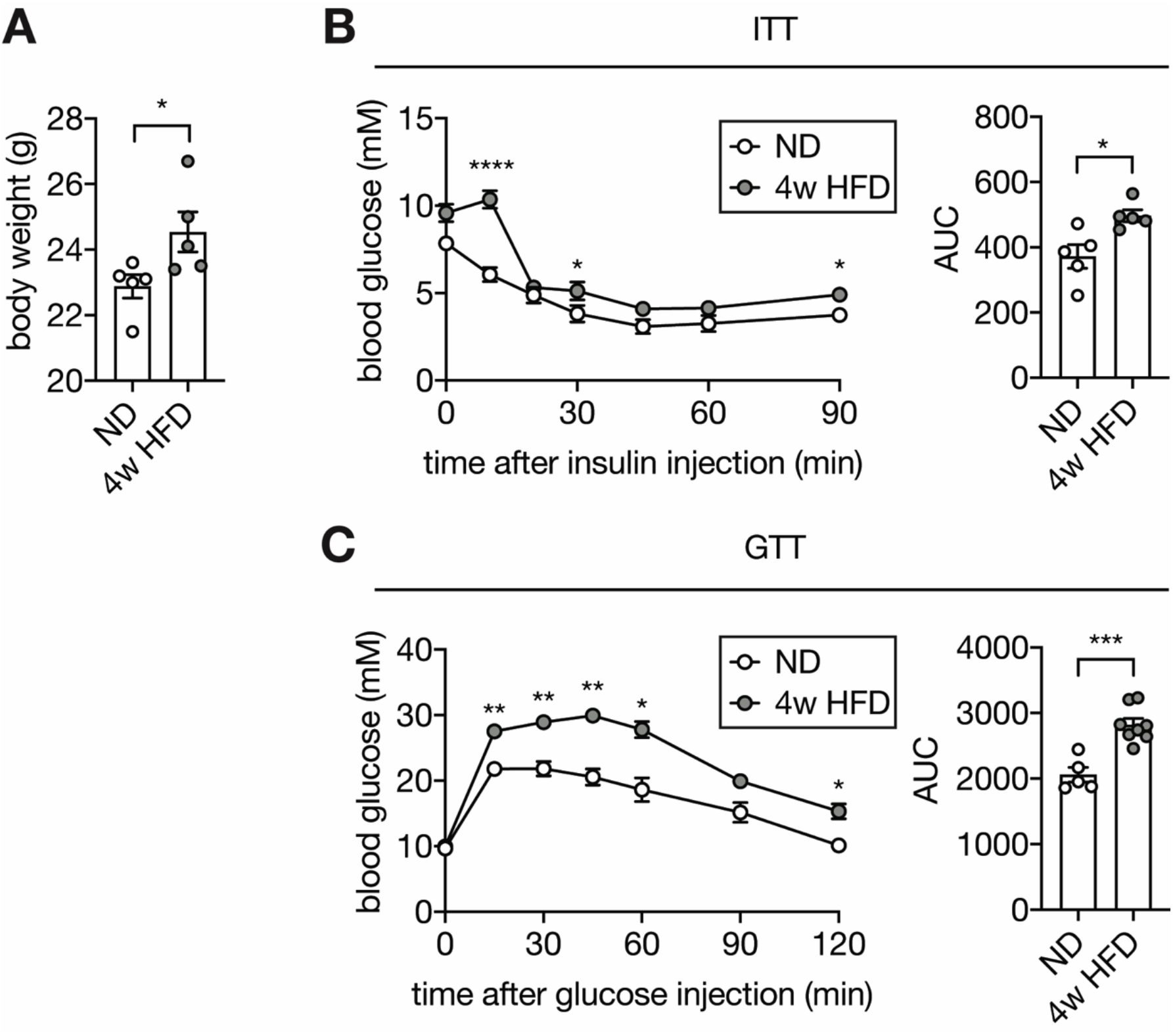
HFD-fed mice are obese, insulin insensitive, and glucose intolerant. (**A**) Body weight of normal diet (ND)- and 4-week high fat diet (HFD)-fed wild-type C57BL/6JRj mice. Student’s t test, *p<0.05. n=5 (ND) and 5 (HFD). (**B**) Insulin tolerance test (ITT) on ND- and 4-week HFD-fed wild-type C57BL/6JRj mice. Mice were fasted for 6 hours and injected with insulin (0.5 U/kg body weight). Two-way ANOVA for glucose curves and Student’s t test for AUC. *p<0.05, ****p<0.0001. n=5 (ND) and 5 (HFD). (**C**) Glucose tolerance test (GTT) on 4-week HFD- and ND-fed wild-type C57BL/6JRj mice. Mice were fasted for 6 hours and i.p. injected with glucose (2 g/kg body weight). Two-way ANOVA for glucose curves and Student’s t test for AUC. *p<0.05, **p<0.01, ***p<0.001. n=5 (ND) and 8 (HFD).

**Fig. S2.**
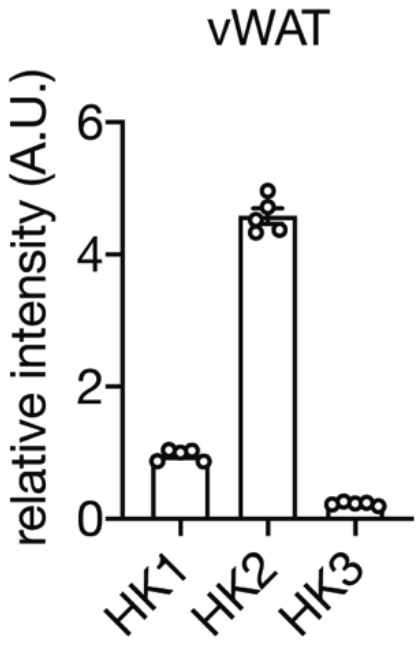
HK2 is the most abundant hexokinase in vWAT. Protein expression of HK1, HK2, and HK3 in vWAT of C57BL/6 mice (Table S1) was normalized to HK1 expression. n=5.

**Fig. S3.**
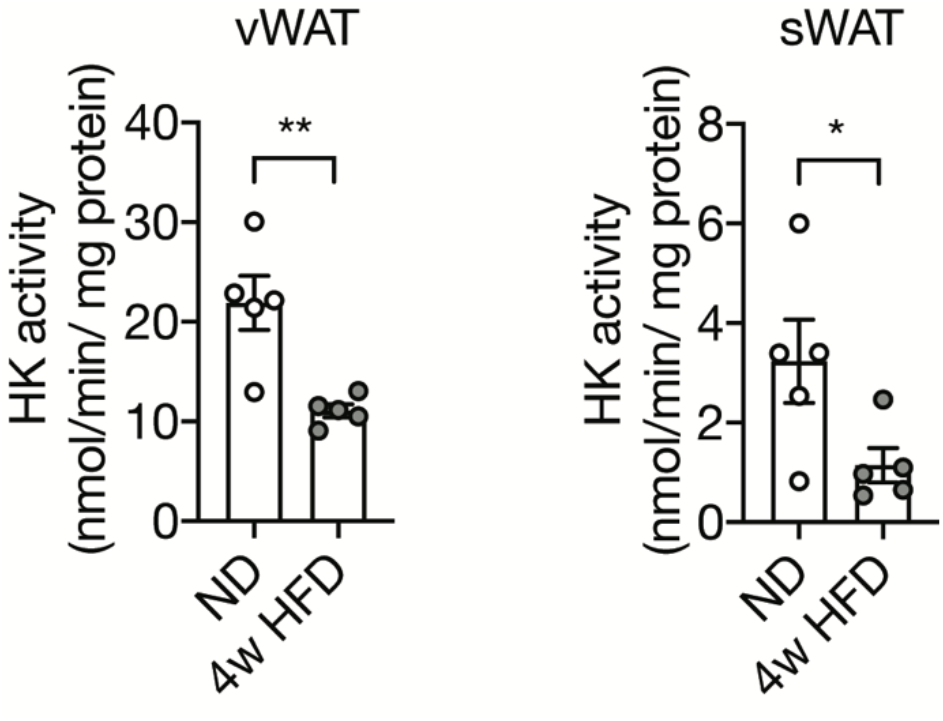
Hexokinase (HK) activity of vWAT and sWAT from ND- and 4-week HFD-fed mice. Student’s t test, *p<0.05, **p<0.01. n=5 (ND) and 5 (HFD).

**Fig. S4.**
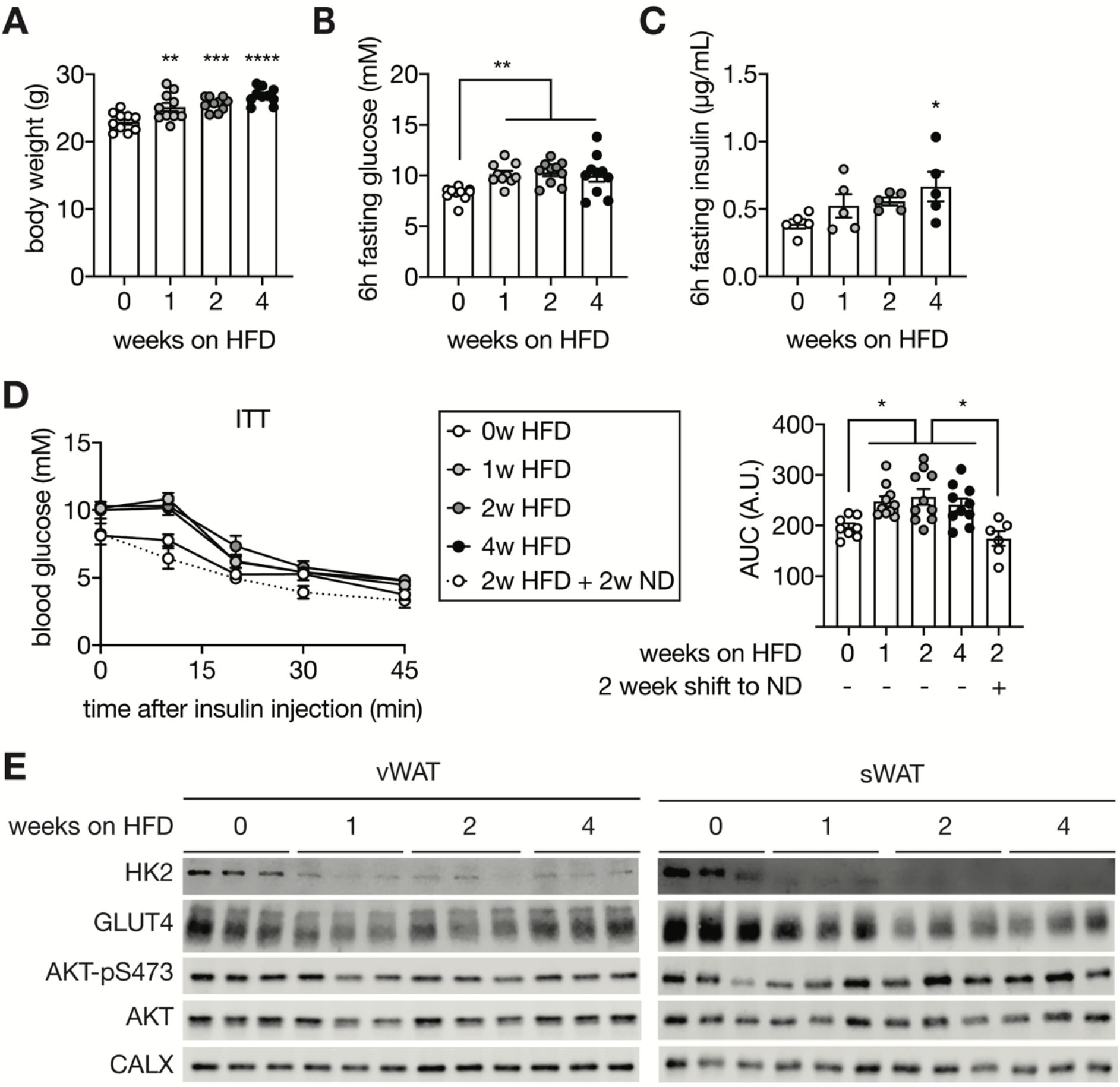
Adipose HK2 expression correlates with hyperglycemia. (**A-B**) Body weight (A) and fasting glucose levels (B) of mice fed a HFD for 0, 1, 2, or 4 weeks. One-way ANOVA compared to 0-week HFD-fed mice, **p<0.01, ***p<0.001, ****p<0.0001. n=10. (**C**) Fasting insulin levels of mice fed an HFD for 0, 1, 2, or 4 weeks. One-way ANOVA compared to 0-week HFD-fed mice, *p<0.05. n=5. (**D**) ITT on mice fed a HFD for 0, 1, 2, 4 weeks, and mice fed 2-week HFD and 2-week ND. Mice were fasted for 6 hours and injected with insulin (0.75 U/kg body weight). Two-way ANOVA, *p<0.05. n=10 (0-week ND), 10 (1-week HFD), 10 (2-week HFD), 10 (2-week HFD), and 6 (2-week HFD + 2-week ND). (**E**) Immunoblot analyses of HK2, GLUT4, AKT-pS473 in vWAT and sWAT from mice fed a HFD for 0, 1, 2, or 4 weeks. Mice were fasted for 6 hours and injected with insulin (0.75 U/kg body weight). n=5.

**Fig. S5.**
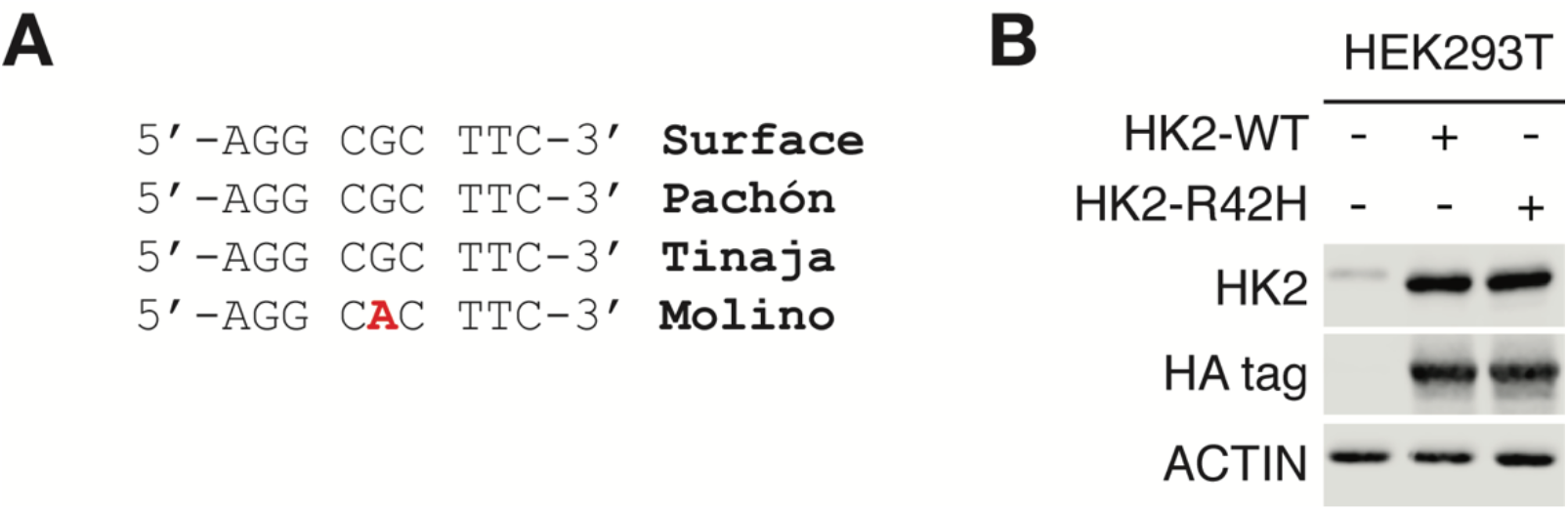
Analyses of R42H mutation. (**A**) DNA sequence of the Molino mutation. (**B**) Immunoblots for lysates of HEK293T cells expressing control, HK2-WT, or HK2-R42H. N=4.

**Fig. S6.**
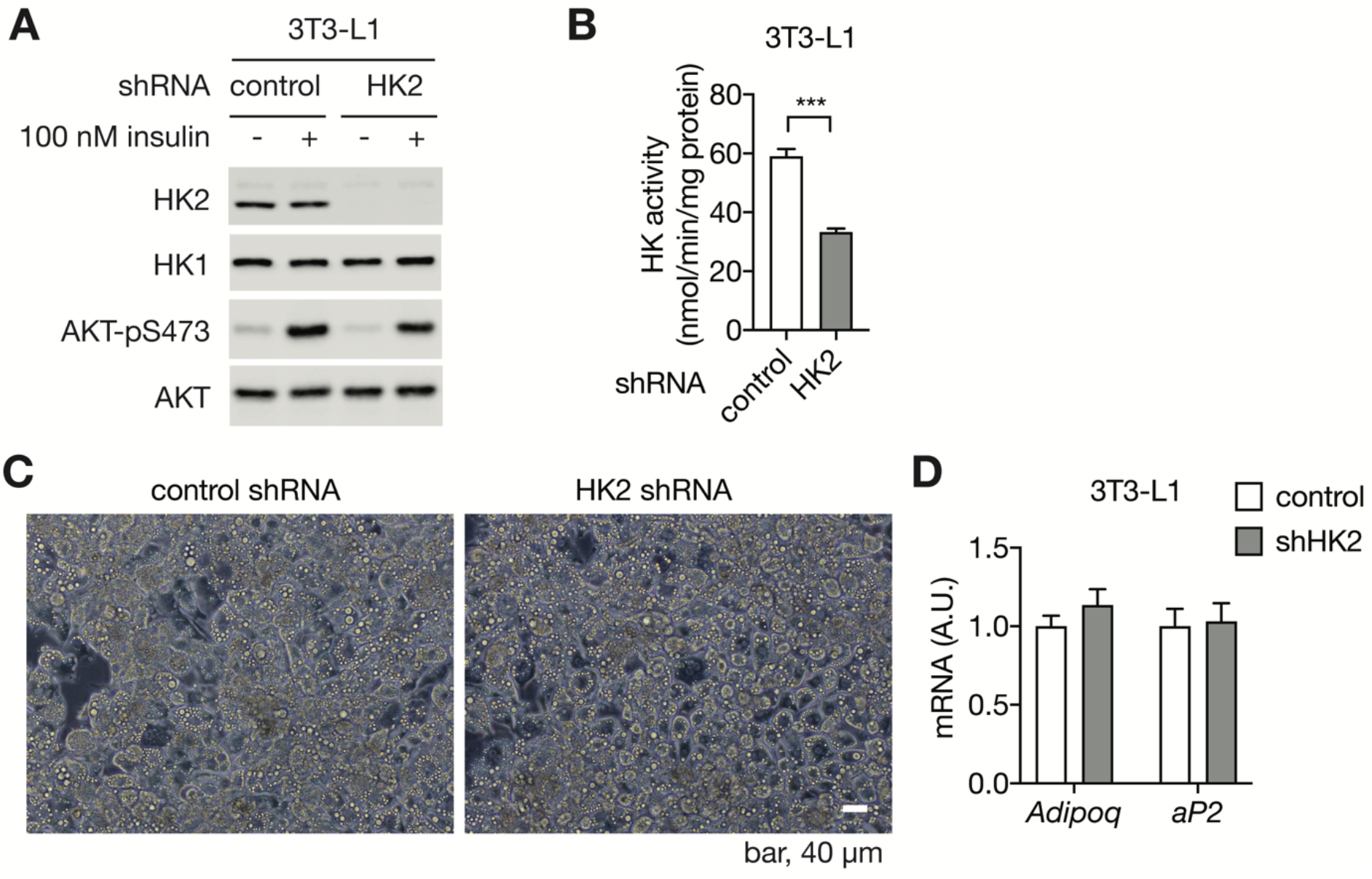
Generation and characterization of HK2 knockdown 3T3-L1 adipocytes. **A.** Immunoblots of control and HK2 knockdown 3T3-L1 adipocytes. Cells were stimulated with 100 nM insulin for 25 min. N=4. **B.** Hexokinase (HK) activity of control and HK2 knockdown 3T3-L1 adipocytes. Student’s t test, ***p<0.001. N=3. **C.** Bright filed images of control and HK2 knockdown 3T3-L1 adipocytes. **D.** mRNA levels of mature adipocyte markers in control and HK2 knockdown 3T3-L1 adipocytes. No significant difference in multiple t test. N=3.

**Fig. S7.**
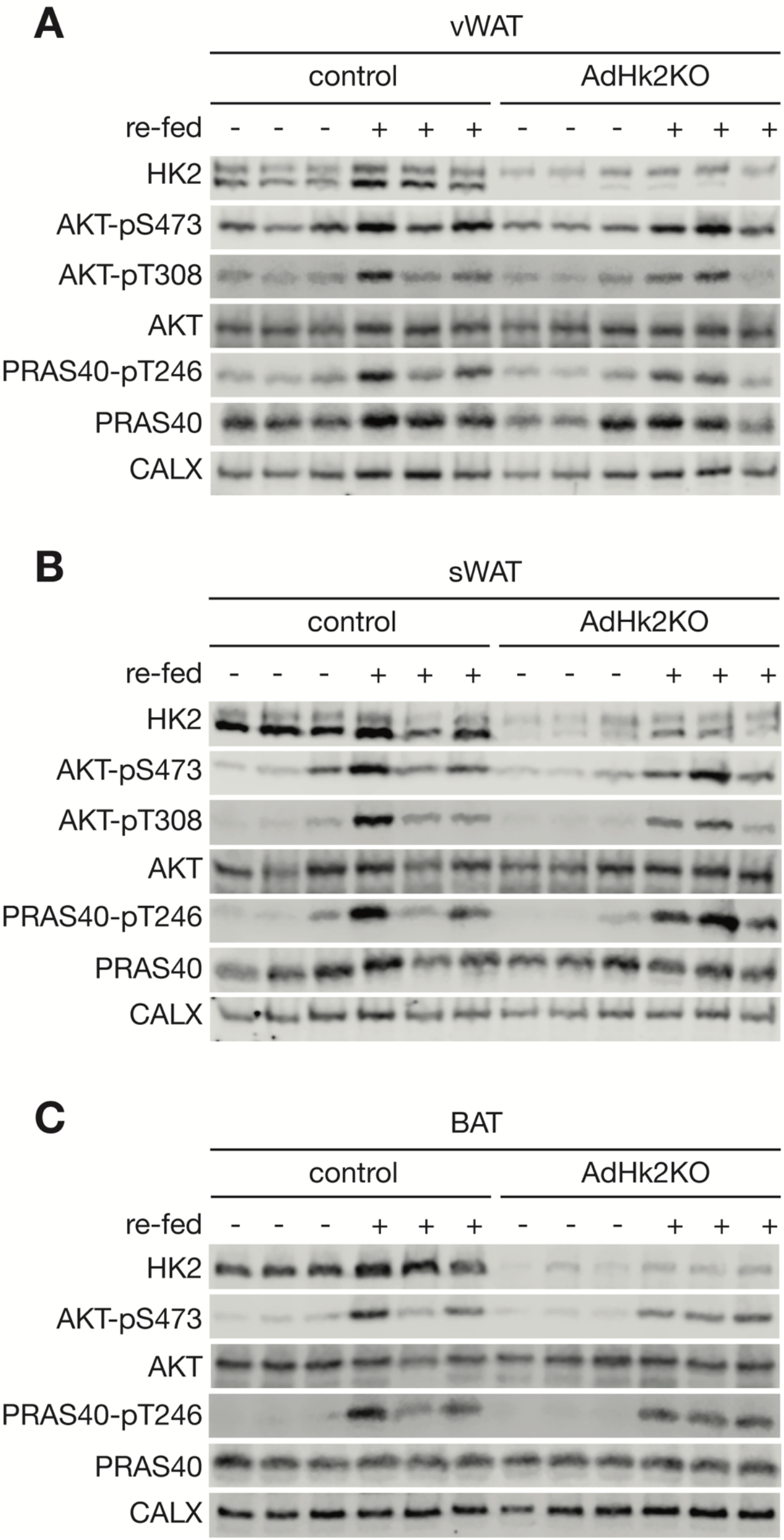
HK2 expression and insulin signaling in adipose tissue of AdHk2KO and control mice. (**A-C**) Immunoblot analyses of HK2 expression and insulin signaling in vWAT (A), sWAT (B), and BAT (C) of control and AdHk2KO mice. Mice were fasted overnight with or without re-feeding for 3 hours. n=6.

**Fig. S8.**
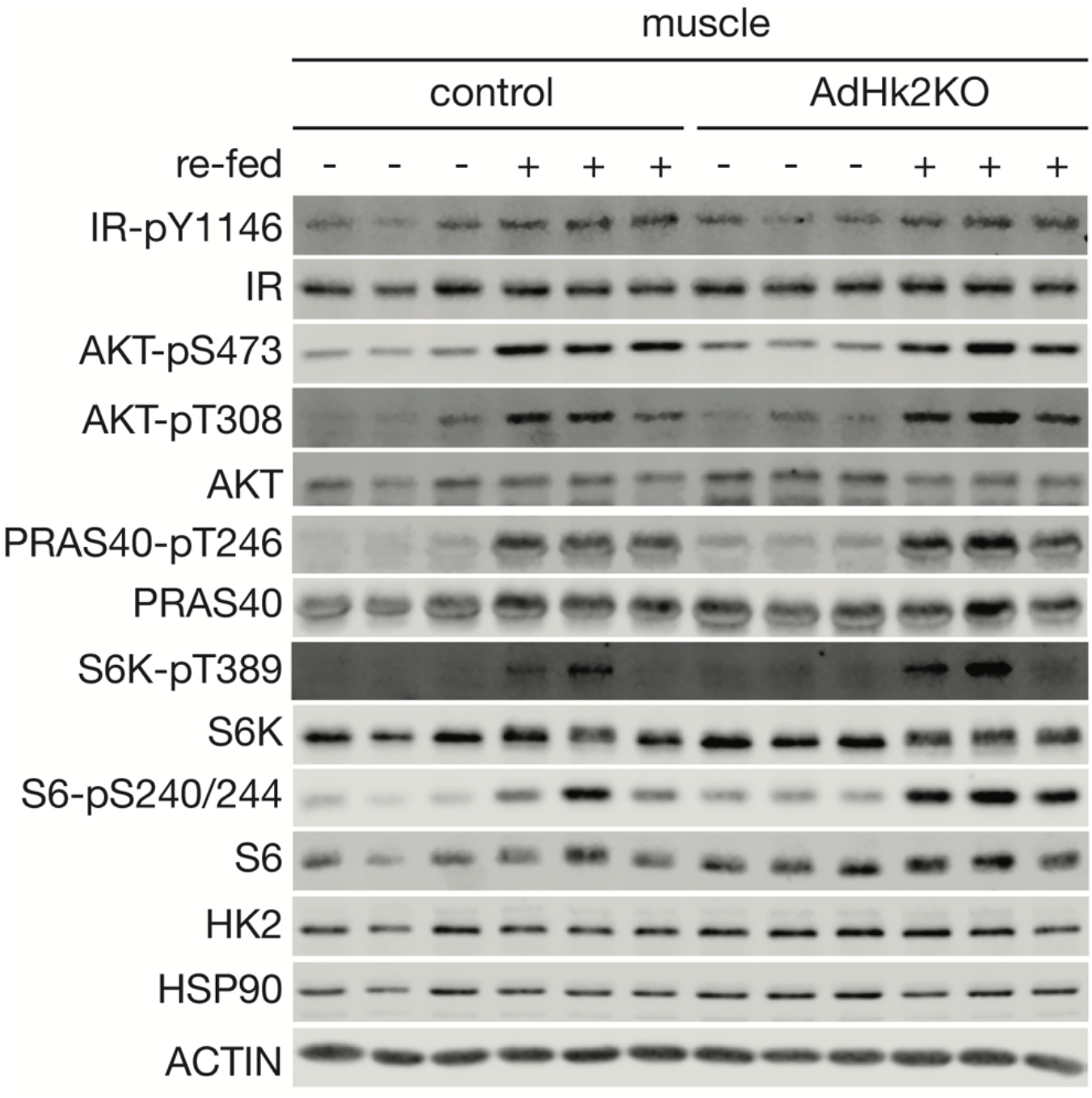
HK2 expression and insulin signaling in skeletal muscle of AdHk2KO and control mice. Immunoblot analyses of HK2 expression and insulin signaling in skeletal muscle of control and AdHk2KO mice. Mice were fasted overnight with or without re-feeding for 3 hours. n=6.

**Fig. S9.**
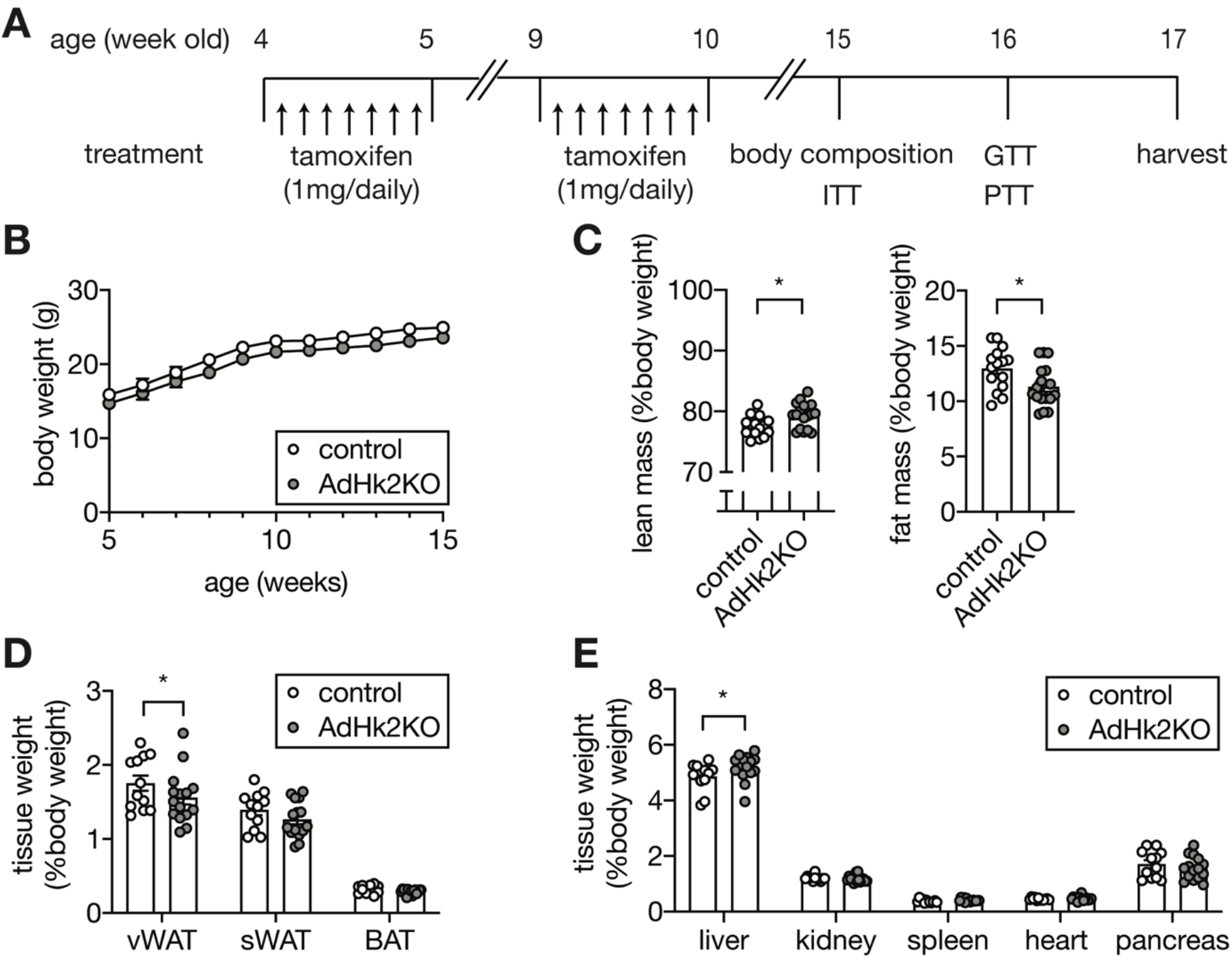
Characterization of AdHk2KO mice. (**A**) Experimental design of tamoxifen treatment, body composition measurement, ITT, GTT, and PTT in control and AdHk2KO mice. Both control and AdHk2KO mice were treated with tamoxifen. (**B-C**) Body weight curve (B), lean and fat mass (C) of control and AdHk2KO mice. Student’s t test, *p<0.05. n=15 (control) and 17 (AdHk2KO). (**D-E**) Organ weight for fat tissues (D) and non-fat tissues (E) of control and AdHk2KO mice. Student’s t test, *p<0.05. n=12 (control) and 15 (AdHk2KO).

**Fig. S10.**
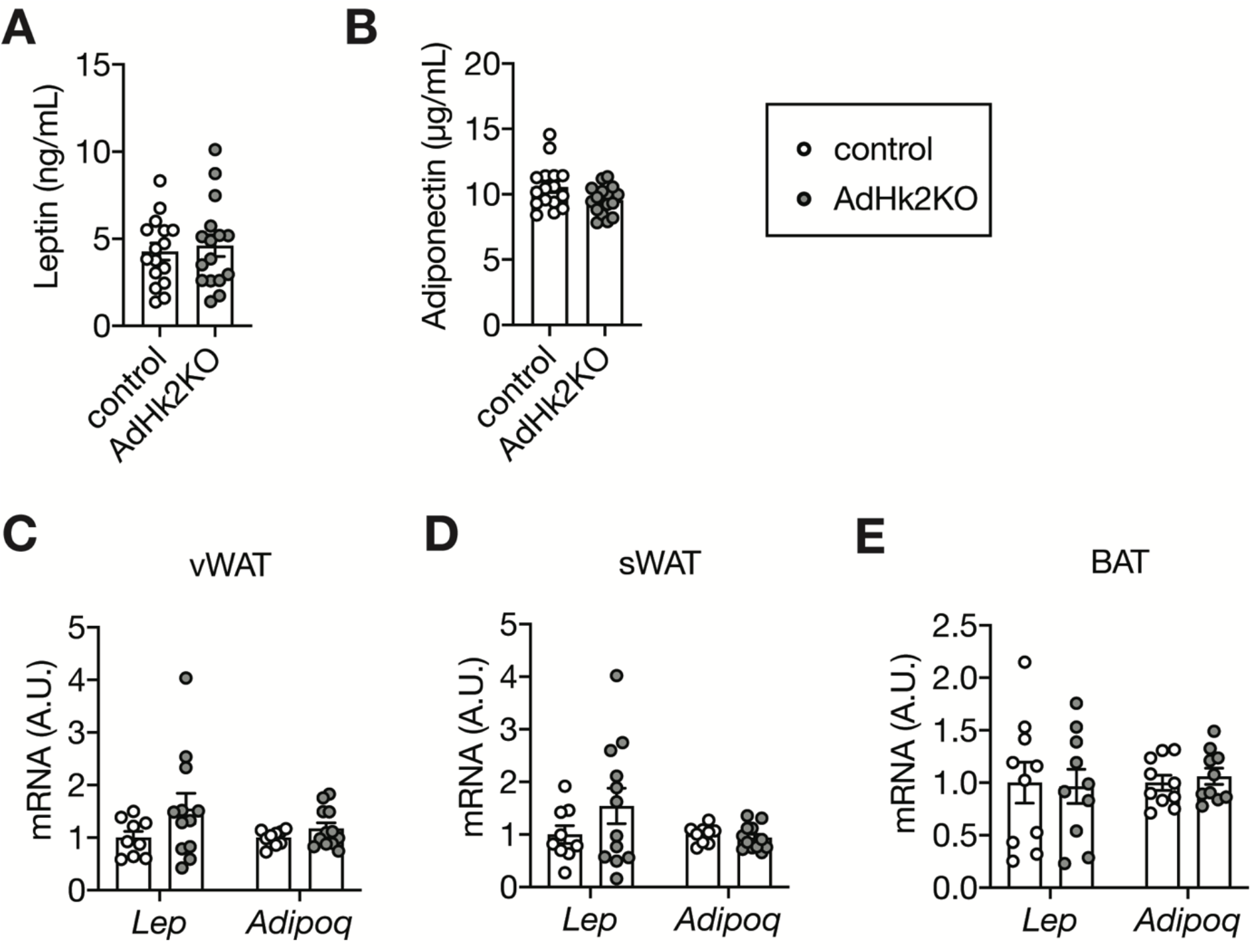
Leptin and Adiponectin levels in control and AdHk2KO mice. (**A**) Plasma leptin levels of ad libitum fed control and AdHk2KO mice. n=16 (control) and 16 (AdHk2KO). (**B**) Plasma adiponectin levels of ad libitum fed control and AdHk2KO mice. n=16 (control) and 16 (AdHk2KO). (**C**) mRNA levels of *Lep* and *Adipoq* in vWAT from control and AdHk2KO mice. n=9 (control) and 12 (AdHk2KO). (**D**) mRNA levels of *Lep* and *Adipoq* in sWAT from control and AdHk2KO mice. n=9 (control) and 12 (AdHk2KO). (**E**) mRNA levels of *Lep* and *Adipoq* in BAT from control and AdHk2KO mice. n=10 (control) and 10 (AdHk2KO).

**Fig. S11.**
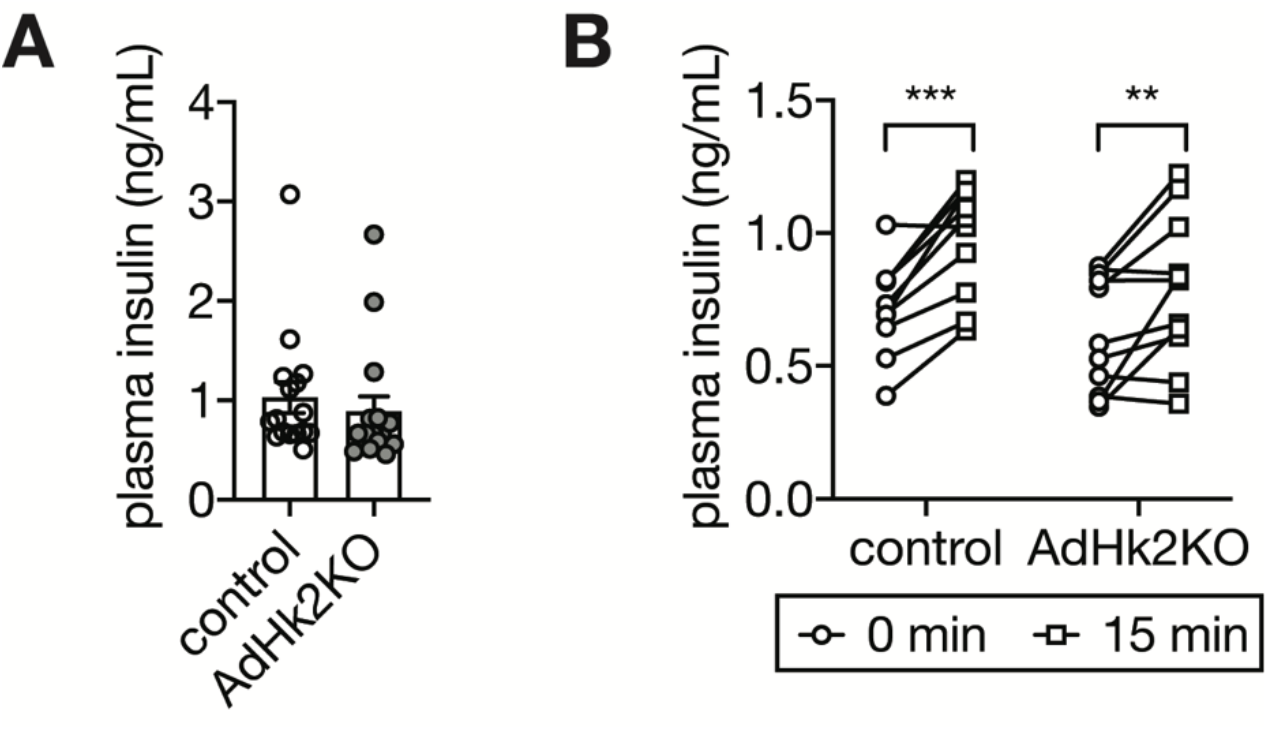
Plasma insulin levels in control and AdHk2KO mice. (**A**) Plasma insulin levels of ad libitum fed control and AdHk2KO mice. n=16 (control) and 16 (AdHk2KO). (**B**) Plasma insulin levels of 6 hour fasted or/and glucose (2g/kg body weight)-injected control and AdHk2KO mice. n=10 (control) and 11 (AdHk2KO).

**Fig. S12.**
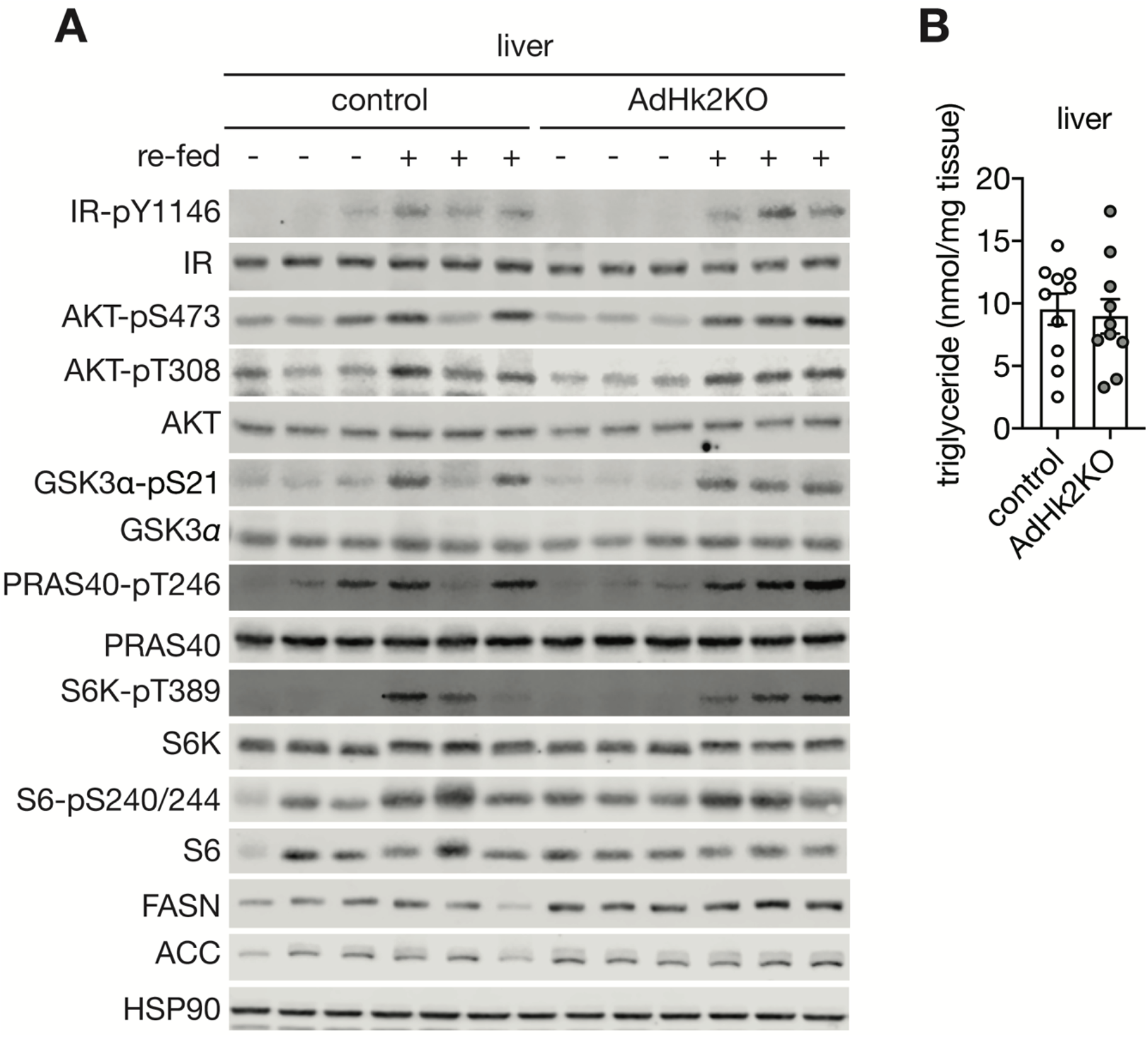
Insulin signaling and triglyceride levels in liver of AdHk2KO and control mice. (**A**) Immunoblot analyses of insulin signaling and lipogenic enzymes in liver of control and AdHk2KO mice. Mice were fasted overnight with or without re-feeding for 3 hours. n=6. (**B**) Hepatic triglyceride levels of control and AdHk2KO mice. n=10 (control) and 10 (AdHk2KO).

**Fig. S13.**
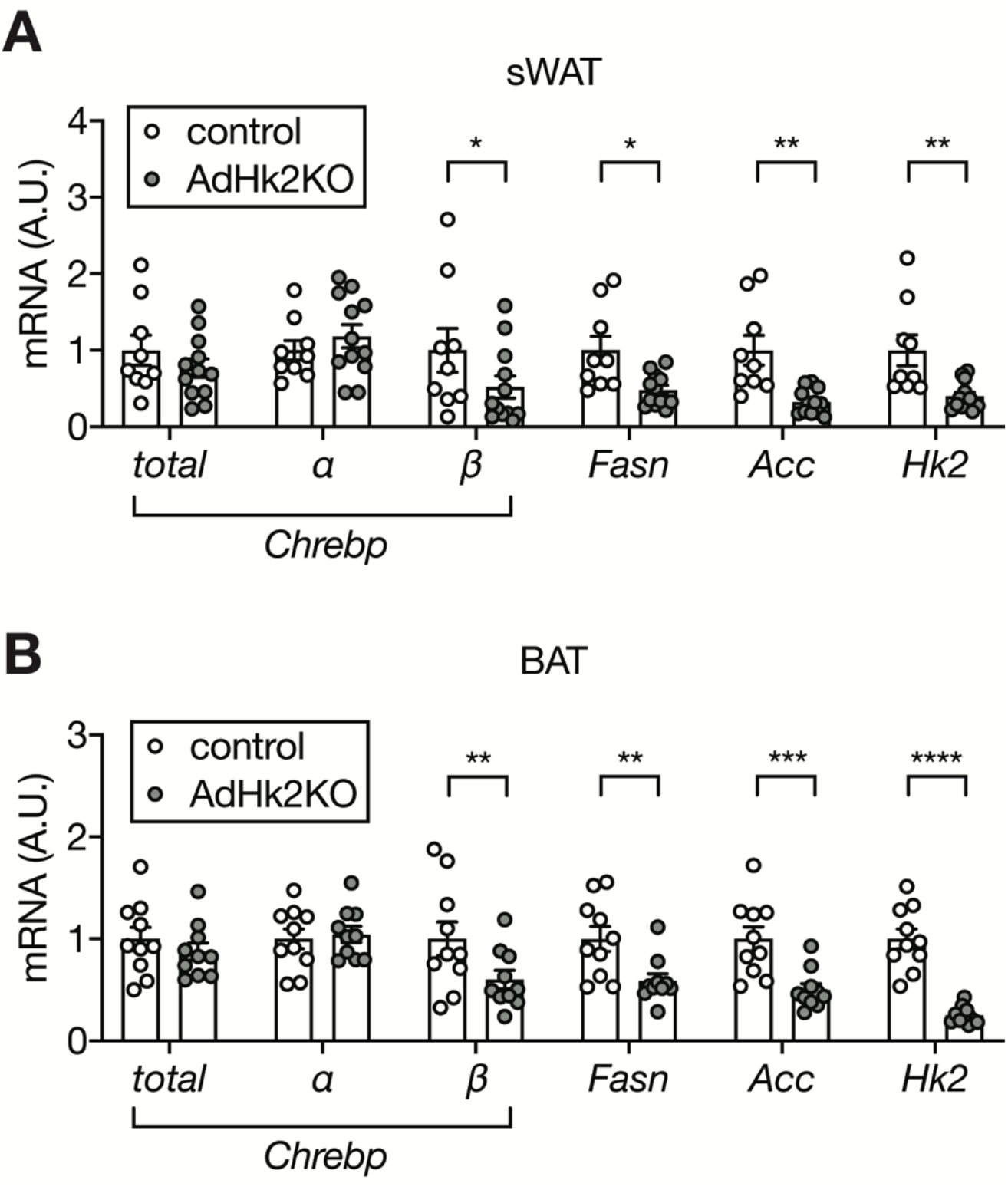
Expression of *Chrebp* and lipogenic genes in sWAT and BAT of control and AdHk2KO mice. (**A**) mRNA levels of lipogenic genes in sWAT from control and AdHk2KO mice. Multiple t test, *p<0.05, **p<0.01. n=9 (control) and 12 (AdHk2KO). (**B**) mRNA levels of lipogenic genes in BAT from control and AdHk2KO mice. Multiple t test, **p<0.01, ***p<0.001, ****p<0.0001. n=10 (control) and 10 (AdHk2KO).

**Fig. S14.**
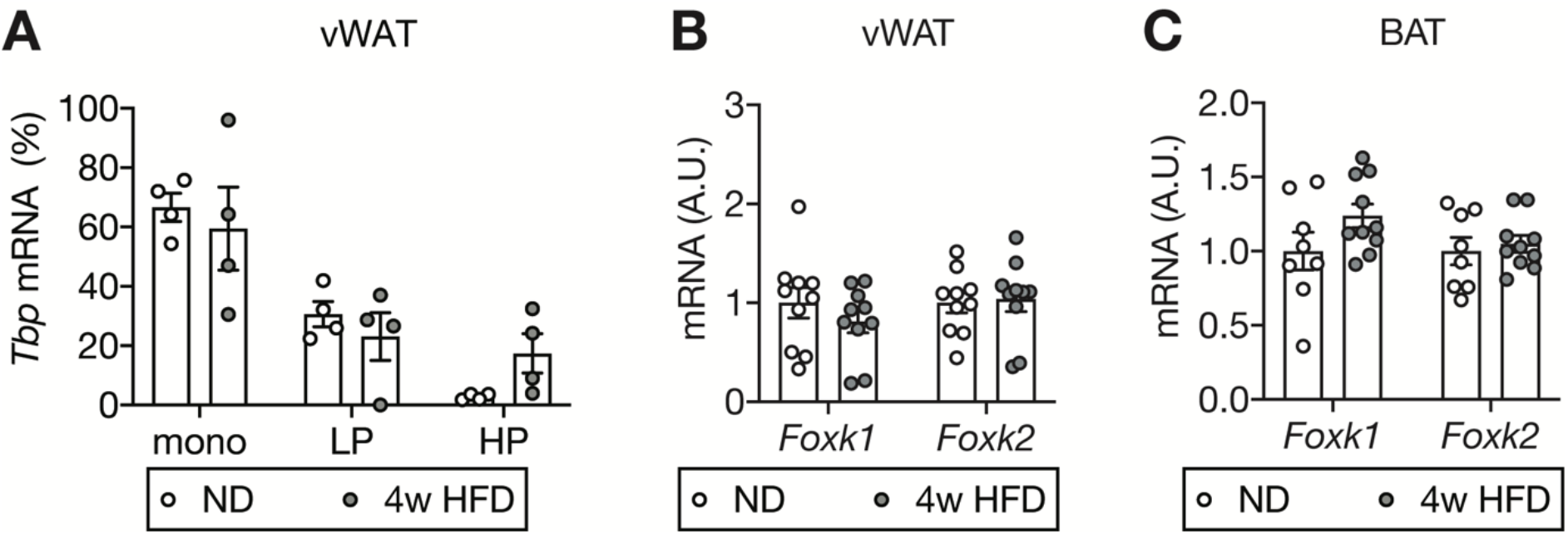
Mechanism of HFD-induced HK2 down-regulation in adipose tissue. (**A**) *Tbp* mRNA abundance in monosome (mono), light polysome (LP), and heavy polysome (HP) fractions in Fig. 4c. No significant difference in two-way ANOVA. n=4. (**B**) *Foxk1* and *Foxk2* mRNA levels in vWAT of ND- or 4-week HFD-fed wild-type C57BL6JRj mice. No significant difference in multiple t test. n=10 (ND) and 10 (HFD). (**C**) *Foxk1* and *Foxk2* mRNA levels in BAT of ND- or 4-week HFD-fed wild-type C57BL6JRj mice. No significant difference in multiple t test. n=8 (ND) and 10 (HFD).

## Tables

**Table S1. Proteome data for vWAT of ND- or 4-week HFD-fed C57BL/6JRj mice.**

**A.** Total proteome in vWAT

**B.** Deregulated proteome in vWAT.

**Table S2. Clinical data for patients.**

